# Interrogating surface versus intracellular transmembrane receptor populations using cell-impermeable SNAP-tag substrates

**DOI:** 10.1101/2020.01.29.924829

**Authors:** Pascal Poc, Vanessa A. Gutzeit, Julia Ast, Joon Lee, Ben J. Jones, Elisa D’Este, Bettina Mathes, David J. Hodson, Joshua Levitz, Johannes Broichhagen

## Abstract

Employing self-labelling protein tags for the attachment of fluorescent dyes has become a routine and powerful technique in optical microscopy to visualize and track fused proteins. However, membrane permeability of the dyes and the associated background signals can interfere with the analysis of extracellular labeling sites. Here we describe a novel approach to improve extracellular labeling by functionalizing the SNAP-tag substrate benzyl guanine (“BG”) with a charged sulfonate (“SBG”). This chemical manipulation improves solubility, reduces non-specific staining and renders the bioconjugation handle impermeable while leaving its cargo untouched. We report SBG-conjugated fluorophores across the visible spectrum, which cleanly label SNAP-fused proteins in the plasma membrane of living cells. We demonstrate the utility of SBG-conjugated fluorophores to interrogate class A, B and C G protein-coupled receptors (GPCRs) using a range of imaging approaches including nanoscopic super-resolution imaging, analysis of GPCR trafficking from intra- and extracellular pools, *in vivo* labelling in mouse brain and analysis of receptor stoichiometry using single molecule pull down.

## Introduction

Membrane receptors, including ligand-gated ion-channels, G protein-coupled receptors (GPCRs), receptor-linked enzymes and, to an extent, transporters, sense extracellular stimuli and convert them into intracellular signals that control cellular function in myriad ways^1^. As such, these proteins are a major focus of drug discovery programs, with GPCRs serving as the largest class of targets^2^. Through an array of approaches, it has become clear that receptor signaling is not restricted to the cell surface but is fine-tuned by a dynamic interplay of receptors both on the surface and in a variety of intracellular compartments^3–6^. Developing techniques for dissecting the relative properties of these distinct pools is an emerging challenge for receptor biology.

Fluorescence microscopy is a powerful technique for direct observation and analysis of molecular processes within a living cell that has been applied extensively to the study of membrane receptors. The continuing development of bright and stable synthetic dyes^7–10^ along with the engineering of self-labelling suicide enzymes, such as SNAP, CLIP and Halo-tags^11^, has spurred the application of targeted, high-resolution imaging in a number of biological contexts^12–21^. Organic dyes that are covalently linked to proteins offer superior brightness, photostability and flexibility compared to fluorescent proteins^10,22^. Many organic dyes are cell permeable and therefore suitable for intracellular labeling. However, this permeability is undesirable when cell surface targeting is required, since confounding background signals can arise from labelled, untrafficked proteins or accumulation of the unlabeled dye in membranes and intracellular compartments. Similarly, fluorescent protein-tagged membrane proteins tend to give high background signals when fused to membrane proteins, since they are expressed, translated and trafficked within the cell. While membrane impermeable fluorophores exist (*e*.*g*. Alexa, Atto or Abberior dyes), many of these fluorophores have been shown to accumulate at the membrane^23^, while the recently developed bright and stable Janelia Fluors^24^ and MaP dyes^8^ are engineered to be membrane permeable. So far, the membrane permeability of a probe has been considered a feature of the fluorophore, with the consequence that imaging only the extracellular protein pool requires changing to a spectrally and photophysically distinct dye as shown recently^25^. While generally useful for qualitative analysis, such an alteration makes quantitative comparisons difficult. Thus, a strategy for rendering dyes impermeable without altering their intrinsic photophysical or spectral properties is needed.

Herein, we describe a subtle, yet powerful modification of *O*^6^-benzylguanine (BG), the substrate for the SNAP-tag, by installing a sulfonate on the leaving group’s C8 position (termed SBG), rendering them impermeable towards the lipid bilayer while conserving reactivity with SNAP. Our general approach allows clean surface labelling of GPCRs in living cells with improved membrane localization and resolution by STED nanoscopy, as well as enhanced signal-to-noise ratio and spread *in vivo*. Moreover, SBG-linked fluorophores open up the possibility to pulse-chase receptors in different compartments, as well as to perform single molecule pulldown (SiMPull) of surface versus intracellular receptor populations. We anticipate that, with this facile strategy, the majority of linked substrates can be rendered impermeable for studies of membrane protein dynamics at the cell surface.

## Results

As a proof-of-principle, we first set out to design and synthesize membrane impermeable versions of SNAP-Cell® TMR-Star and SNAP-Cell® 647-SiR, two popular commercially-available fluorophores for SNAP-tag labeling (**Fig. 1a; Scheme S1**). Based on previous studies, which report that alterations at guanine position C8 are tolerated for enzymatic SNAP labelling^26^, we hypothesized that substituents on the BG would alter the permeability of the entire compound without interfering with labeling. Conveniently, this moiety would also be liberated upon SNAP labeling, thus removing any potential alterations to the photophysical properties of the protein bound fluorophore itself. As such, three moieties were examined and prepared as their TMR- and SiR-bearing reagent, namely the parent BG (with H at C8: “BG-TMR” and “BG-SiR”), a previously described^26^ carboxylate CBG (with a linked COOH at C8: “CBG-TMR” and “CBG-SiR”) and, finally a sulfonate (with a linked SO_3_H at C8: “SBG-TMR” and “SBG-SiR”) (**Fig. 1b, c**). Bearing in mind that sulfonates display a *pK*_*a*_ < 0, SBG will be permanently negatively charged in physiological buffers, thereby unable to cross the lipid bilayer membrane and, presumably, repelled further by the negatively charged surface. Accordingly, SBG-TMR and SBG-SiR were obtained by coupling sulfonate-containing taurine to a protected CBG precursor, which after deprotection allowed straightforward amide coupling to NHS-activated fluorophores (see SI).

**Figure 1:**
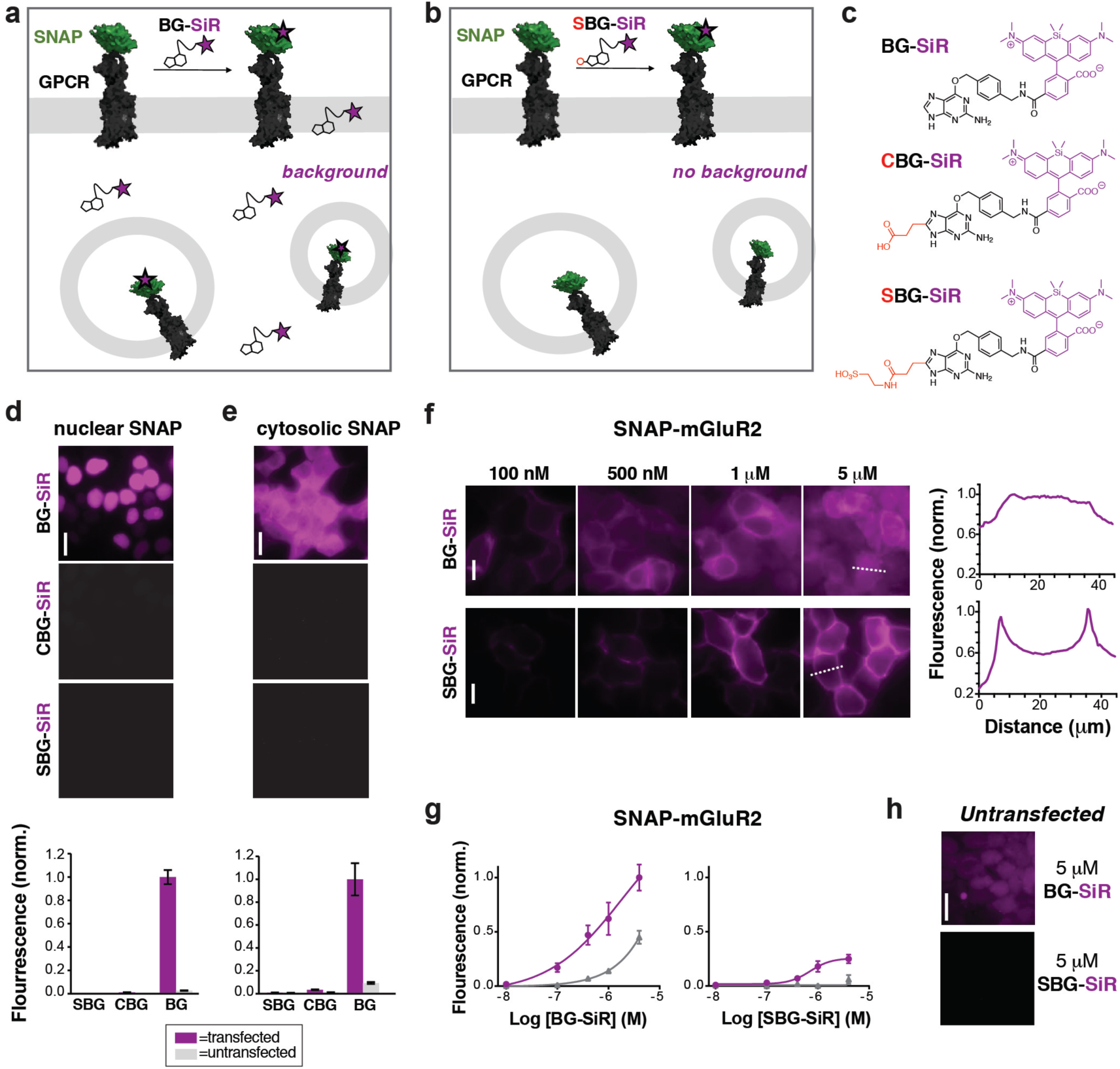
CBG and SBG-conjugated SiR are membrane-impermeable and enable specific targeting of surface proteins. **a**) Application of permeable BG-SiR to N-terminally SNAP-fused GPCRs leads to extracellular labelling of surface receptors and background signals due to labelling of intracellular pools and non-specific dye accumulation. **b**) Application of impermeable SBG-SiR should lead solely to labelling of extracellular tags with reduced background. **c**) Chemical structures of BG-SiR, CBG-SiR and SBG-SiR. **d, e**) BG-SiR, but not CBG-SiR and SBG-SiR, labels nucleus-targeted (**d**) or cytosol-targeted (**e**) SNAP-tags. **f, g**) Concentration-dependent labelling of SNAP-mGluR2 leads to intracellular background signals using BG-SiR, which is absent using SBG-SiR. Line scans, right, demonstrate that labeling restricted to the surface only with SBG-SiR. **h**) Untransfected cells show background signals from BG-SiR but not from SBG-SiR. Scale bars are 20 µm.

For initial assessment of labelling properties we chose TMR since covalent binding of a non-fluorogenic dye can easily be observed using fluorescence polarization. As expected, BG-TMR, CBG-TMR and SBG-TMR showed complete SNAP labeling *in vitro* as assessed by full protein mass spectrometry^27^ (**Fig. S1-S4**) with labelling kinetics ∼3-times slower for SBG-TMR (*t*_1/2_ = 51.3 seconds) *versus* BG-TMR (*t*_1/2_ = 17.8 seconds) yet complete within minutes (**Fig. S5a**). An advantage of using a charged residue is increased solubility, and as such, SBG-TMR can be readily dissolved in pure PBS at a concentration >1.5 mM, while BG- and CBG-TMR need to be dissolved in DMSO (>1 mM) before dilution in PBS for usage. More importantly, BG-TMR (∼80 µM in PBS, 1% DMSO) was not stable in solution at room temperature, precipitating within minutes to leave a steady-state concentration of ∼17 µM in the supernatant (**Fig. S5b**). Notably, SBG-TMR remained in solution at ∼70 µM without the addition of DMSO over three days.

We next tested the ability of modified BGs to label intracellular SNAP by expressing either a cytosolic- or nuclear-targeted SNAP-tag before applying BG, CBG or SBG-conjugated fluorophores. While labeling with 500 nM BG-SiR for 45 minutes at 37 °C produced clear fluorescence for both cytosolic and nuclear SNAP-tags, labeling with 500 nM CBG-SiR or SBG-SiR did not produce any substantial signal with either construct in transiently transfected HEK293 cells (**Fig. 1d, e**). Notably, background fluorescence in untransfected cells was highest for BG-SiR, lower for CBG-SiR and undetectable for SBG-SiR (**Fig. 1d, e**). As such, we decided to continue our characterization with SBG-SiR because it showed a robust decrease in membrane permeability compared to CBG-SiR. We next labeled cells expressing a SNAP-tagged GPCR, metabotropic glutamate receptor 2 (“SNAP-mGluR2”), with either BG-SiR or SBG-SiR. Both compounds produced clear fluorescence over a similar range of labeling concentrations (**Fig. 1f-h**), but signals from SBG-SiR were more confined to the plasma membrane (**Fig. 1f, right**) and showed less background labeling in untransfected cells (**Fig. 1g, h**). Together, these data validate the idea that addition of an anionic sulfonate group to BG can render an attached fluorophore membrane-impermeable for targeting of surface proteins with reduced non-specific labeling.

Due to their distinct spectral and photophysical properties, different fluorophores are required for multimodal applications. Based on the desire to prevent membrane permeability of different fluorophores at will, we asked if this approach was generalizable to a family of fluorophores spanning visible to far-red wavelengths. To do this, we synthesized and tested SBG-conjugated Oregon Green (OG), TMR, Janelia Fluor 549 (JF_549_) and Janelia Fluor 646 (JF_646_) (**Fig. 2a-d**), the latter showing superior brightness and photostability over their tetramethyl and silicon rhodamine counterparts^24^. In all cases, SBG-conjugated fluorophores clearly label surface receptors, with minimal labeling of intracellular SNAP-tags (**Fig. 2a-d**).

**Figure 2:**
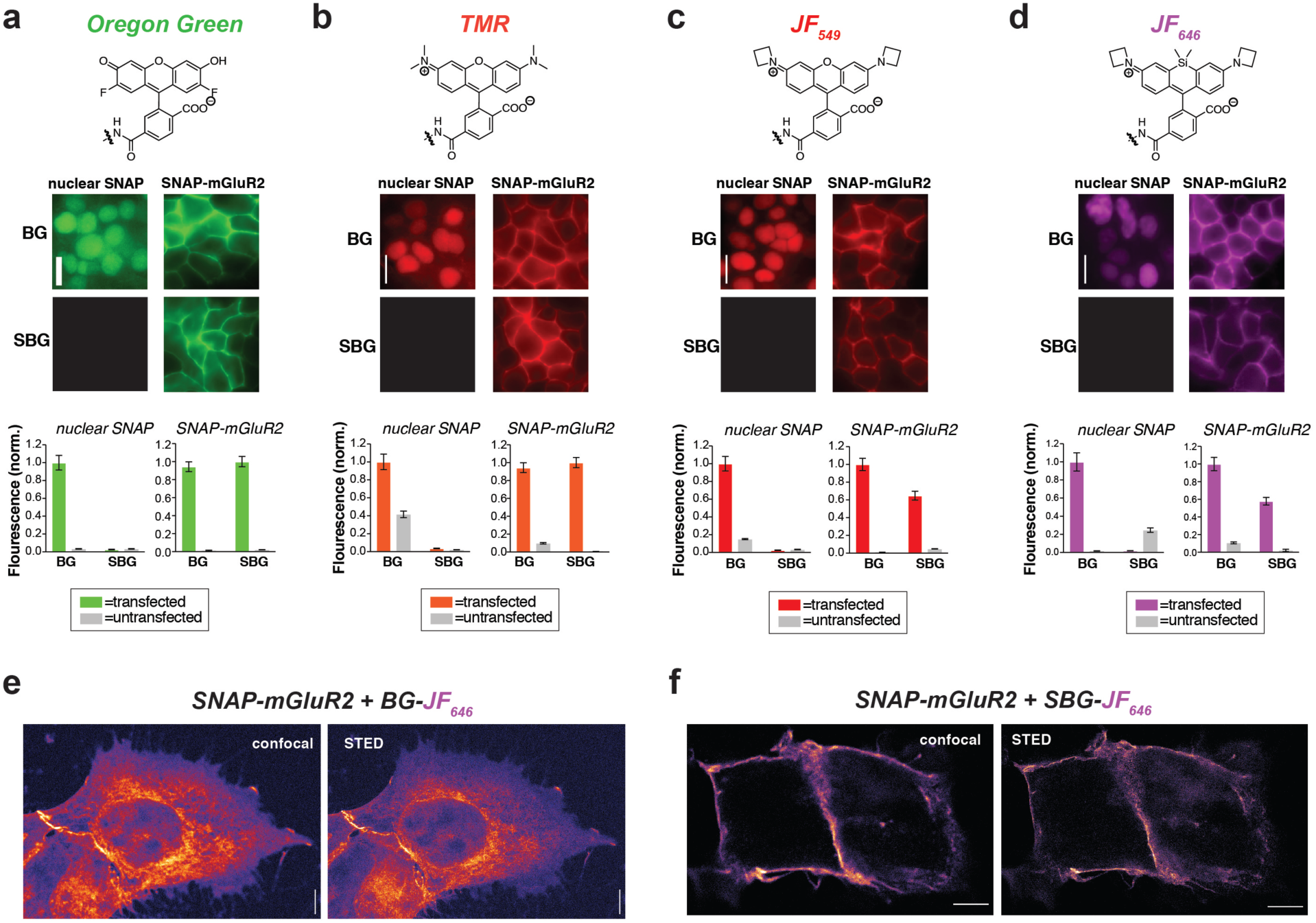
SBG-conjugated fluorophores across the visible spectrum enable surface-specific SNAP labeling and nanoscopic imaging of surface receptors. **a-d**) SBG conjugation enables surface targeting of Oregon Green (**a**), TMR (**b**), JF_549_ (**c**), and JF_646_ (**d**). All fluorophores are able to label nuclear SNAP-tags when conjugated BG but not SBG and show cleaner surface targeting of SNAP-mGluR2. **e, f**) Confocal and superresolution STED nanoscopy of mGluR2 using BG-JF_646_ (**e**) and SBG-JF_646_ (**f**) shows clear isolation of the membrane population only using the impermeable SBG probe. Data is represented as mean ± SEM. Scale bars are 20 µm.

To test if membrane-localized SNAP-tag labels offer advantages for cell biology, we turned to nanoscopic STED imaging. A dye with outstanding far-red performance in STED microscopy with respect to photostability and brightness is JF_646_^28^. Accordingly, we used JF_646_ SNAP-tag probes to target SNAP-mGluR2 in transiently transfected HEK 293 cells. Similar to what was observed by widefield microscopy (**Fig. 2d**), we observed mainly intracellular staining in fixed cells with BG-JF_646_ (**Fig. 2e**). This intracellular fluorescence is likely due to a mixture of immature proteins that have not yet trafficked to the cell surface and surface receptors that have been internalized. By instead using SBG-JF_646_, we obtained images where the dye remained solely at the cell surface, and furthermore, were able to resolve membranes with a lateral resolution of 91±23 nm using STED nanoscopy (n=42; *cf*. FWHM_confocal_ = 295±85 nm, n=35) (**Fig. 2f**).

We next asked if SBG-conjugated fluorophores could allow for superior labelling of GPCRs *in vivo*. We recently established SNAP-tag labeling *in vivo* in the frontal cortex of living mice using local injection of BG-conjugated fluorophores^29^. Presumably, the high solubility and reduced cell permeability of SBG-conjugated fluorophores should lead to improved tissue staining. Based on our prior study, we virally-delivered SNAP-mGluR2 into the medial prefrontal cortex (mPFC) of adult mice before injecting BG-JF_549_ or SBG-JF_549_ 8 weeks later at the same coordinates (**Fig. 3a**). Clear labeling was observed with both compounds (**Fig. 3b-g**), but we observed a larger spread of SBG-JF_549_ fluorescence in transduced brains (**Fig. 3d; Fig. S6**) and untransduced brains showed a 2-fold higher background for BG-JF_549_ than its SBG counterpart (**Fig. 3g**).

**Figure 3:**
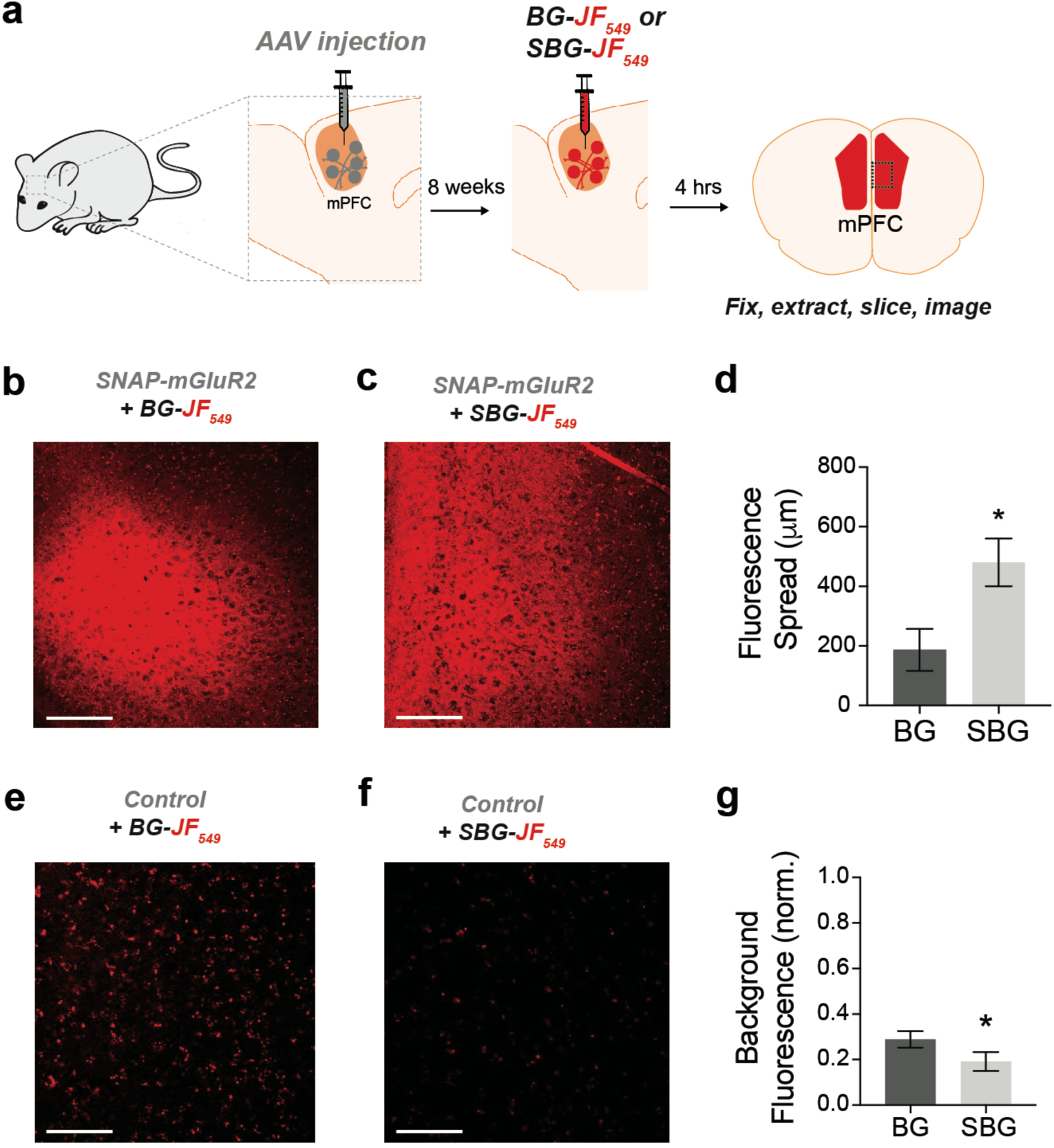
*In vivo* labeling of a SNAP-tagged receptor with SBG-conjugated fluorophores produces less background and more spread. **a**) Schematic showing AAV-mediated expression of SNAP-mGluR2 in the medial prefrontal cortex of mice, followed by SBG-JF_549_ or BG-JF_549_ dye injection and slice preparation 8 weeks later. **b-c**) Representative images showing fluorescence in slices from SNAP-mGluR2 expressing mice following injection of BG (**b**) or SBG (**c**) fluorophores. **d**) SBG-JF_549_ shows broader spread throughout the cortex compared to BG-JF_549_. * indicates statistical significance (unpaired t-test, p=0.04). **e, f**) Representative images showing fluorescence in control slices following injection of BG (**e**) or SBG (**f**) fluorophores. **g**) Larger background signals are observed for BG-conjugated dye. * indicates statistical significance (unpaired t-test, p=0.007). Data is represented as mean ± SEM and comes from n=3 mice for each condition. Scale bars are 150 µm.

Having established efficient surface-targeted labeling with SBG-conjugated fluorophores, we next asked if we could use BG- and SBG-conjugated fluorophores to separate the respective intra- and extracellular pools of a membrane receptor. We employed glucagon-like peptide-1 receptor N-terminally fused to SNAP (“SNAP-GLP1R”), and used two spectrally separated dyes to pulse-chase label different receptor pools (**Fig. 4a**). GLP1R is involved in glucose homeostasis^30^ and is known to undergo rapid endocytosis and trafficking upon activation with the agonist Exendin4(1-39) (Ex4). Live tracking of surface-exposed receptors has previously been achieved by using BG-Alexa or BG-Atto dyes^31,32^ or by the use of specific antibodies and fixation at given time points^30,32^. However, we wanted to test the ability to simultaneously analyze the surface pool and the intracellular pool that has been translated but not yet trafficked to the membrane. As such, we incubated cells with SBG-TMR (500 nM) to label SNAP-GLP1R at the cell surface, followed by BG-SiR (500 nM) to label the intracellular, residual and newly trafficked (*i*.*e*. during the wash step) surface receptor pools (**Fig. 4a**). After washing, SNAP-GLP1R was clearly labelled at the surface with TMR and intracellularly with SiR (**Fig. 4b**). No bleedthrough was apparent in controls that used only one dye (**Fig. S7**). Surface TMR:SNAP-GLP1R was activated by Exendin4(1-39) (Ex4; 25 nM), before tracking of TMR and SiR-labelled receptor pools in live cells using high-resolution confocal microscopy (**Fig. 4b**). After 15 minutes of agonist treatment, cells were washed and the antagonist Exendin4(9-39) (Ex4; 500 nM) was applied to allow the internalized and cytosolic receptors pools to be sorted and re-trafficked to the surface (**Fig. 4b**). As expected, TMR-labelled GLP1R readily internalized following ligand binding, before trafficking and degradation upon washout and application of antagonist (**Fig. 4c**), as evaluated by mean fluorescence intensity at the membrane and in the cytosol (**Fig. 4c, d**). Interestingly, we noticed a cytosolic pool of SiR:SNAP-GLP1R, which either remained static and did not undergo trafficking, or alternatively, was degraded before being replenished by the portion of surface receptor labelled by SiR (**Fig. 4c, d**). Thus, GLP1R present at intracellular sites immediately before orthosteric activation are unlikely to contribute to ligand-induced receptor turnover.

**Figure 4:**
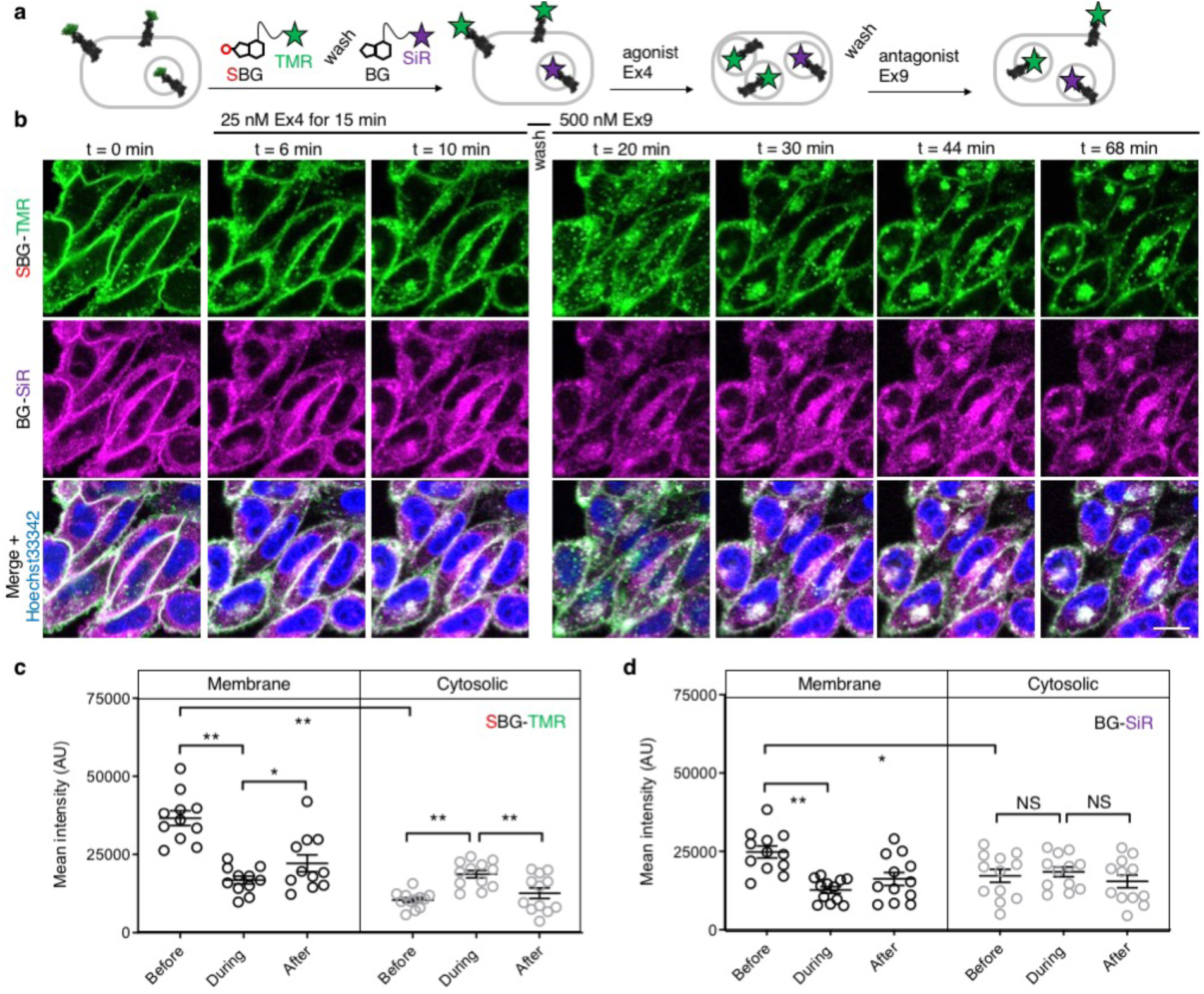
SBG and BG-conjugated fluorophores allow tracking of different receptor pools in live cells. **a**) Surface GLP1R is labelled with SBG-TMR, before washing and labeling of cytosolic receptor (and residual or newly trafficked receptor) with BG-SiR. The two GLP1R pools are then tracked over time in response to agonist stimulation (Ex4, Exendin4(1-39); 25-50 nM), followed by washing and antagonist application (Ex9, Exendin9(9-39); 500 nM) to halt trafficking. **b**) Cytosolic GLP1R (BG-SiR) remains relatively static, while surface GLP1R (SBG-TMR) reversibly internalizes (representative images shown) (scale bar = 34 µm) (nuclei are labelled with Hoechst33342). (**c, d**) Quantification of mean fluorescence intensity at the membrane and within the cell, showing a significant increase in cytosolic SBG-TMR (**c**), but not BG-SiR (**d**), signal before (0 min), during (11-17 mins) and after (53-61 min)- agonist stimulation (repeated measures two-way ANOVA, Fishers LSD or Bonferonni’s post-hoc test; F = 10.93, DF = 2) (n = at least 2 different imaging positions in 6-9 wells, 3 independent repeats). *P<0.05, **P<0.01, NS, non-significant.

We next wondered if our technique could be used to probe the stoichiometry of GPCR populations inside the cell *versus* at the plasma membrane. Fluorescence-based methods have been widely used for assessing GPCR dimer- and oligomerization but rarely distinguish between surface and intracellular pools which may lead to confounding results and discrepancies across studies. This is especially critical as GPCR homo- and hetero-multimerization remains a controversial topic that may have major implications for general physiology and drug discovery^33^. We decided to use our labeling probes in conjunction with single molecule pulldown (“SiMPull”) a strategy which allows single receptor complexes to be isolated and imaged for analysis of stoichiometry *via* counting of fluorophore bleaching steps^34^. To probe a prototypical class C GPCR, reported to form constitutive dimers by most studies to date^35,36^, we used HA-SNAP-mGluR2. Conversely, an HA-SNAP-beta-2 adrenergic receptor construct (“HA-SNAP-ß2AR”) was used as a prototypical class A GPCR, which has been found as a monomer or dimer or higher order oligomer depending on experimental conditions^34,37–41^. Each construct was labeled with either SBG-JF_549_, to label only surface receptors, or SNAP-Surface® Block followed by BG-JF_549_ (see methods for details) to isolate intracellular receptors. Cell imaging showed distinct fluorescence patterns for each receptor depending on the labeling paradigm (**Fig. 5a**) and labeling controls indicated that the BG-surface block prevented >95% of labeling of surface receptors without altering the efficiency of labeling intracellular receptors (**Fig. S8**). Following labeling, cells were lysed and detergent-solubilized GPCRs were isolated for single molecule imaging at a low density on passivated coverslips using an anti-HA antibody as previously described^42^ (**Fig. 5b**). Single molecules were imaged using TIRF microscopy to allow for stepwise fluorophore bleaching which could be used to measure receptor stoichiometry (**Fig. 5c**). SBG-JF_549_ labeled HA-SNAP-mGluR2 showed ∼55-60% 2-step photobleaching, consistent with previous studies indicating the formation of strict mGluR dimers^35,36,42^. However, intracellular receptors labeled with BG-JF_549_ showed reduced 2-step bleaching, indicating reduced dimerization in this population (**Fig. 5d**). These data suggest that a portion of the intracellular receptors are immature and monomeric.

**Figure 5:**
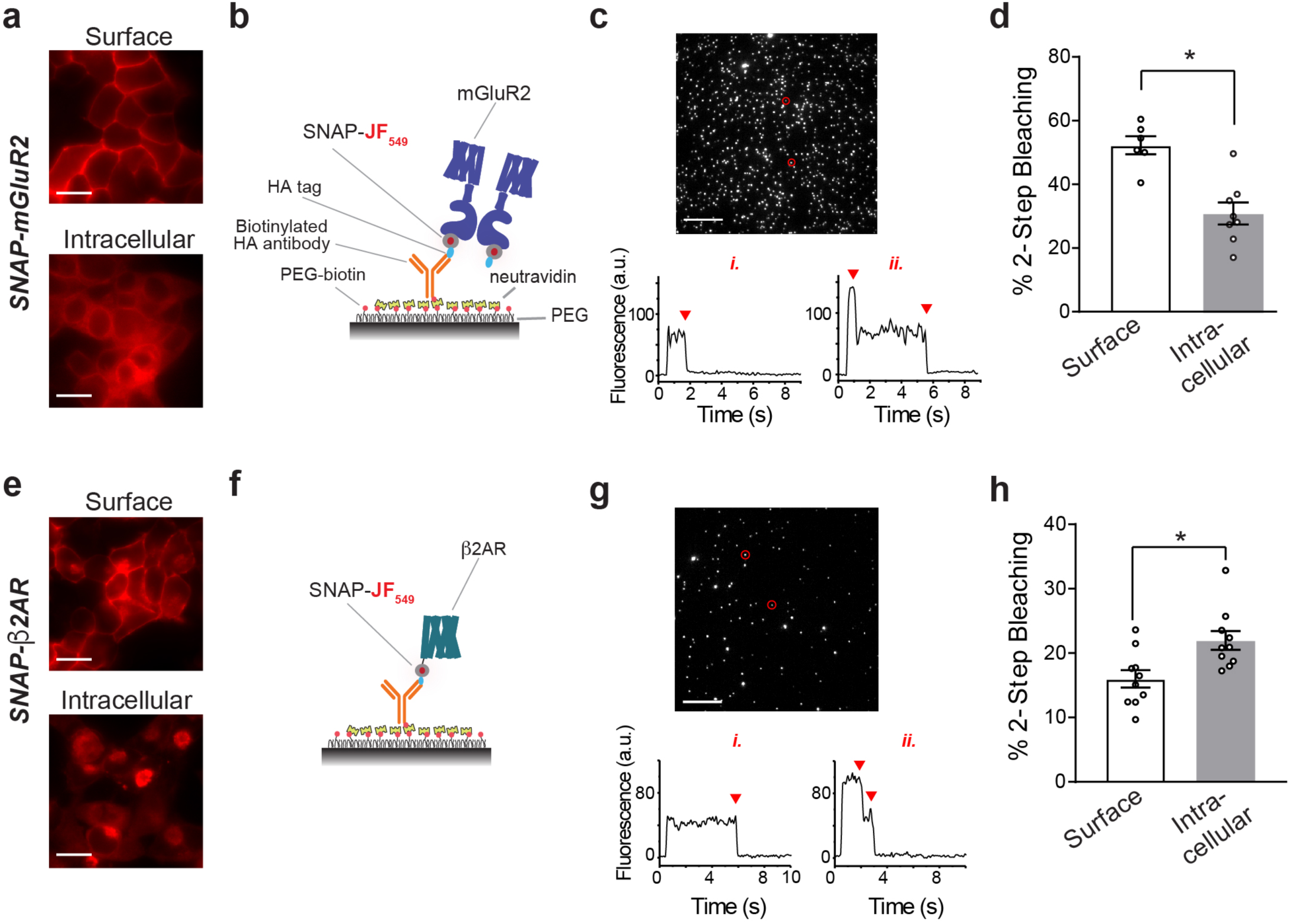
BG and SBG-conjugated fluorophores enable SiMPull analysis of isolated surface or intracellular receptor populations. **a**) Representative images showing labeling of either surface (top) or intracellular (bottom) SNAP-mGluR2 with SBG- or BG-JF_549_, respectively. **b**) Schematic showing single molecule pulldown configuration where an anti-HA antibody is used to isolate a sparse surface of SNAP-tagged mGluR2 following fluorophore labeling. **c**) Representative image of single molecules for SNAP-mGluR2, with representative bleaching traces for a 1-step and 2-step example (bottom). Note: >95% of spots bleached in either 1 or 2-steps. **d**) Summary of the proportion of 2-step bleaching steps for each labeling configuration. Each point represents one independent movie and bars show mean ± SEM. * indicates statistical significance (unpaired t test, p=0.0005). **e-h**) Same as **a-d** but with SNAP-ß2AR. * indicates statistical significance (unpaired t test, p=0.008). Scale bars are 10 µm.

We next performed the same experiment with HA-SNAP-ß2AR (**Fig. 5e, f**). Consistent with our previous SiMPull study^42^, we found weak dimerization of surface receptors labeled with SBG-JF_549_ (**Fig. 5g, h**). However, when we targeted intracellular receptors a small, but significant increase in apparent dimerization was observed (**Fig. 5h**). Together these data demonstrate the suitability of SBG and BG dyes for isolating surface *versus* intracellular receptor pools for experiments that take place *in vitro* following cell lysis. In addition, they indicate that different receptor pools may have different distributions of monomeric and multimeric receptors, emphasizing the importance of identifying which pool is being probed in a given study.

Taken together, we report the design and use of novel highly-soluble and membrane impermeable probes for the examination of different GPCRs from the purified protein to live cells to the whole organism.

## Discussion

The use of SNAP-tags to specifically label and interrogate membrane-spanning proteins has proven a powerful technique in cell biology. To further improve this method, we have rationally designed a modification to the BG leaving group to yield sulfonated-BG (SBG) that renders a range of *a priori* permeable fluorophores impermeable towards the plasma membrane. As such, the fluorophore remains identical after labelling, without alteration of its spectral or other photophysical properties. It should be noted that derivatizing the leaving group is possible for nucleobases as used for SNAP, but not for the leaving group of the Halo-tag, being a chloride atom. Recent approaches have used charged moieties synthetically introduced between the leaving group and the dye^43^, with the need to test for influences on binding kinetics and fluorogenicity, the latter which is optimized for the protein surface it is exposed to. Another approach is the use of inherently impermeable dyes, such as some Alexa, ATTO or Abberior, which display properties different to the dyes we aimed to use. Other impermeable modifications, such as relatively large quenchers custom tailored for the fluorophore, have been reported for no-wash labelling of charged fluorophores^44^. In contrast, we describe a minimal alteration, independent of the cargo that should be generally applicable.

Using our cell-impermeable SNAP substrates, we showcase fast and clean membrane staining of SNAP-mGluR2, accompanied by STED nanoscopy. By using SBG-linked bright and photostable dyes, we could restrict labelling to the lipid bilayer for different dyes in the visible spectrum. We furthermore obtained highly resolved images of SNAP-mGluR2 residing at the membrane using STED in fixed cells, which proved to be impossible after using the BG-version due to high intracellular background staining. The most stable and widely used far-red STED dyes (Atto 647N, STAR RED and STAR635/P) are neither membrane permeable nor fluorogenic and hence cannot be used in cases where a comparison between intracellular and extracellular ligands is needed. On the other hand, the best performing dyes for live STED imaging (SiR, JF_646_, CP 610) have been designed to ensure membrane permeability^9,24,45^ and, therefore, they also do not allow for a comparison between intracellular and membrane protein pools, which necessarily requires the use of SBG-ligands.

JF_549_ showed superior behavior when applied as its SBG-version *in vivo*. After application via injection, SBG-JF_549_ showed a 2-fold reduction in background when compared to its BG-congener. In contrast, its spread in virally-infected brains was markedly increased. These results, in addition to the ability to solubilize dyes without a co-solvent (*i*.*e*. DMSO), demonstrate the power of our simple chemical modification for use in living animals.

We were also able to stain a SNAP-GLP1R fusion construct at the membrane with SBG-TMR and the remaining, mostly intracellular, pool with BG-SiR. Separating pools of the same protein has been achieved before, for instance by using fluorogen activated protein (FAP)^46,47^ probes or the fluorescence-activating and absorbance-shifting tag (FAST)^48^. While these previous studies rely on non-covalent labelled protein tags fused to the BK channel or a transmembrane helix, respectively, we report a SNAP-fusion to a blockbuster targeted GPCR. In addition, our system does not rely on Förster Resonance Energy Transfer (FRET), which adds another layer of complexity and need for additional control experiments, as has been shown for malachite green conjugates. Furthermore, our approach allows for the use of different colors in the same experiment, while the FAST system uses charged and non-charged forms of the same fluorophores. As such, we show that only GLP1R present at the membrane before ligand stimulation undergoes trafficking in response to activation. GLP1R which is already present inside the cell prior to stimulation never reaches the membrane. As such, two pools of GLP1R likely exist in the unstimulated state: 1) surface-exposed receptor which is trafficking-competent in the presence of ligand; and 2) internalized, cytoplasmic, newly-synthesized or incorrectly processed GLP1R which slowly traffics to the membrane in the absence of ligand or is, alternatively, degraded and replenished by labelled residual membrane receptor. Since peptide ligand cannot enter the cell, it is unlikely that the internalized GLP1R pool contributes meaningfully to intracellular (*e*.*g*. endosomal) signaling responses. What is the relevance of such observations for GLP1R function? Firstly, the same GLP1R pool might constantly turnover with continuous ligand stimulation, with the intracellular pool never making it to the membrane within the imaging timescale used here (*i*.*e*. measuring dynamic changes after activation). Secondly, ligands that specifically target the intracellular GLP1R pool might further increase efficacy of GLP1R agonists used in the treatment of metabolic disease. As such, SBG-fluorophores as well as other SBG-conjugated species are likely to open up interesting avenues of investigation for both GLP1R biology and the broader study of protein trafficking.

Finally, we also demonstrate the value of the SBG approach for chemically-tagging surface receptors for subsequent biochemical isolation. We use this to show that SBG-targeted surface GPCRs can display different stoichiometries than BG-targeted intracellular GPCRs. In the case of the class C GPCR mGluR2 intracellular receptors, presumably immature proteins, we were able to show reduced dimerization compared to the strict dimerization of the cell surface population. This result suggests that dimeric receptors are preferentially trafficked to or maintained on the cell surface. In contrast, intracellular ß2AR showed enhanced dimerization compared to surface pools. Critically, the ability to use the same fluorophore (JF_549_) for each condition, removes any possibility that differences in photobleaching pattern is due to differences in dye photophysics. Future work will be needed to dissect the determinants of the differential dimerization of these populations, their sensitivity to different stimuli and to assess this phenomenon across a range of GPCRs and other membrane proteins. The flexible control afforded by SBG-conjugated fluorophores will be critical for such studies.

In conclusion, we report the design and use of novel highly-soluble and membrane impermeable probes for the interrogation of different GPCRs from the purified protein level to live cells to the whole organism.

## Online Methods

### Synthesis

Chemical synthesis (Supporting Scheme 1, 2) and characterization of compounds is outlined in the Supporting Information. All compounds were >95% purity according to HPLC if not stated otherwise.

### SNAP_f_ expression, purification, and mass spectrometry after labelling

SNAP_f_ was expressed and purified as described previously.^27^ Briefly, SNAP_f_ with an N-terminal Strep-tag and C-terminal 10xHis-tag was cloned into a pET51b(+) expression vector for bacterial expression and complete amino acid sequences for constructs used can be found in the Supporting Information. For purification, SNAP_f_ was expressed in the *E. coli* strain BL21 pLysS. LB media contained ampicillin (100 µg/mL) for protein expression. A culture was grown at 37 °C until an OD_600_ of 0.6 was reached at which point cells were induced with IPTG (0.5 mM). Protein constructs were expressed overnight at 16 °C. Cells were harvested by centrifugation and sonicated to produce cell lysates. The lysate was cleared by centrifugation and purified by Ni-NTA resin (Thermofisher) and Strep-Tactin II resin (IBA) according to the manufacturer’s protocols. Purified protein samples were stored in 50 mM HEPES, 50 mM NaCl (pH 7.3) and either flash frozen and stored at −80 °C. For SNAP_f_ labelling, 25 µL of 30 µM dye (BG-/CBG-/SBG-TMR) in activity buffer (50 mM HEPES, 50 mM NaCl, pH = 7.3) was added to a 10 μM solution of SNAP_f_ in activity buffer in 1.5 mL Eppendorf tubes. This resulted in a 3-fold excess of labelling substrate and mixing was ensured by carefully pipetting the solution up and down. The reaction mixture was allowed to incubate at r.t. for 1 h before tubes were stored at 4 °C until MS analysis.

### SNAP_f_ labelling kinetics

Kinetic measurements were performed on a TECAN Spark 20M platereader by means of fluorescence polarization. Stocks of SNAP_f_ (400 nM) and TMR substrates (100 nM) were prepared in activity buffer: 50 mM HEPES, 50 mM NaCl, 1 mM DTT, 100 ng/mL BSA, pH = 7.3. SNAP_f_ and substrates were mixed (50 μL each) in a Greiner black flat bottom 96 well plate and fluorescence polarization reading was started immediately (λ_Ex_ = 535±25 nm; λ_Em_ = 595±35 nm; 30 flashes; 40 µs integration time). Experiments were run in triplicates, data was normalized and one-phase decay fitted in GraphPad Prism 7.

### Solubility studies

Lyophilized compounds were dissolved in PBS (SBG-TMR) or in DMSO (BG-TMR). Concentration was assessed by diluting each 1:50 into PBS/0.1% SDS and measuring UV absorbance at 550 nm by a NanoDrop (extinction coefficient: 90,000 M^-1^ cm^-1^; pathlength d = 0.1 cm) to be in the single digit millimolar range. BG-TMR was diluted 1:100 in PBS and aliquoted into 1.5 mL Eppendorf tubes, which were spun at 15,000 rpm for 30 seconds, the supernatant diluted 1:1 with PBS/0.1% SDS and concentration determined at a NanoDrop. Time intervals were 0, 7, 14, 21 and 40 minutes. SBG-TMR was diluted 1:25 in PBS and aliquoted into 1.5 mL Eppendorf tubes, which were spun at 14,600 rpm for 30 seconds, the supernatant diluted 1:1 with PBS/0.1% SDS and concentration determined at a NanoDrop. Time intervals were 0, 7, 14, 21, 40 minutes and after 3 days (4320 minutes).

### Expression and fluorescence imaging in HEK 293T cells

HEK 293T cells were cultured in DMEM with 5% FBS, seeded on 18 mm poly-L-lysine-coated cover slips in a 12-well plate and transfected using Lipofectamine 2000 (Thermo Scientific). Cells were transfected with 0.3-0.7 µg/well of SNAP-tagged constructs.

After 24-48 h of expression, cells were first washed with extracellular (EX) solution containing (in mM): 10 HEPES, 135 NaCl, 5.4 KCl, 2 CaCl_2_, 1 MgCl_2_, pH 7.4 and labeled with fluorophores at 37°C at the indicated concentrations for 45 minutes. An inverted microscope (Olympus IX83) was used for fluorescence imaging. Live cell images were captured using a 60x objective (NA 1.49) with an exposure time of 100 ms. Laser intensity was kept constant across the compared samples for fluorescence intensity quantification. Average fluorescence intensity from cell images was measured using ImageJ by drawing a region of interest (ROI) around cell clusters. Fluorescence intensity values from multiple images were then averaged. Each condition was tested in at least 2 separate transfections.

### Super-resolution microscopy

HEK293 cells transfected with SNAP-mGluR2 growing on 18 mm coverslips were treated with 400 nM BG-JF_646_ or SBG-JF_646_ for 60 minutes in DMEM (without phenol red), washed and fixed (4% paraformaldehyde for 20 min, followed by quenching solution 0.1 M glycine, 0.1 M NH4Cl in PBS). Cells were mounted in mowiol supplemented with DABCO and imaged on an Abberior STED 775/595/RESOLFT QUAD scanning microscope (Abberior Instruments GmbH, Germany) equipped with STED lines at λ = 595 and λ = 775 nm, excitation lines at λ = 355 nm, 405 nm, 485 nm, 561 nm, and 640 nm, spectral detection, and a UPlanSApo 100×/1.4 oil immersion objective lens. Following excitation at λ = 640 nm, fluorescence was acquired in the spectral window λ = 650-800 nm. FWHM was measured on raw data and calculated using Fiji software with Gaussian fitting.

### Mice

All animal use procedures were performed in accordance with Weill Cornell Medicine Institution Animal Care & Use Committee (IACUC) guidelines under approved protocol (2017-0023). Male wild-type mice were of strain C57BL/6J and purchased from Jackson Laboratory and were maintained under specific pathogen free conditions at the Weill Cornell Medicine Animal Facility. Animals were provided food and water ad libitum and housed in a temperature and humidity controlled environment with a 12 hour light/12 hour dark cycle.

### *in vivo* SNAP labeling

AAV-mediated expression of SNAP-mGluR2 and in vivo SNAP labeling was performed as previously described^29^. Briefly, Male C57BL/6J mice were injected at p60 with a 1:1 viral cocktail of AAV9-EF1a-FLEX-SNAP-mGluR2-WPRE-hGH (Penn Vector Core) and pENN-AAV9-CamKII 0.4-Cre-SV40 (Addgene) or, as a control, only AAV9-EF1a-FLEX-SNAPmGluR2-WPRE-hGH. Mice were injected in the medial prefrontal cortex (AP + 1.85, ML +/- 0.35, DV-2.2, −1.8) with 500 nL per site using a Kopf stereotaxic and World Precision Instruments microinjection syringe pump with a 10 µL syringe and 33g blunt needle. 8 weeks after viral injection, mice received infusion of 500 nL of 1 µM BG-JF_549_ or SBG-JF_549_ targeted to the same site as viral injection. 4 hours later mice underwent transcardial perfusion and were fresh fixed with 4% paraformaldehyde. Brains were extracted and bathed in 4% paraformaldehyde for 24 hours followed by 72 hours in 30% sucrose PBS solution. Brains were mounted and frozen at −20 °C in OCT block and medial prefrontal cortex was sliced at 40 µm thick on a cryostat at −22 °C. Slices were wet mounted to glass slides and secured with coverslip using VECTASHIELD HardSet Antifade Mounting Medium with DAPI (Vector Laboratories). Glass slides were imaged using an Olympus Confocal FV3000 and images were processed and analyzed in ImageJ.

### GLP1R trafficking studies

CHO-K1 cells stably expressing the human SNAP-GLP1R (Cisbio) (CHO-K1_SNAP-GLP1R) were maintained in DMEM supplemented with 10% FCS, 1% penicillin/streptomycin, 500 µg/mL G418, 25 mM HEPES, 1% nonessential amino acids and 2% *L*-glutamine. Cells were incubated with 500 nM SBG-TMR for 15 minutes at 37 °C, 5% CO2, before washing three times in medium. BG-SiR was then applied at 500 nM for 20 minutes under the same conditions. Live imaging was performed using a Zeiss LSM880 meta-confocal microscope configured with GaAsP detectors, a 63×/1.2 W Korr M27 objective and a temperature and CO2-controlled chamber. Exendin4(1-39) (25-50 nM) and Exendin4(9-39) (500 nM) were applied at the indicated timepoints and concentrations. SBG-TMR, BG-SiR and Hoechst33342 were excited using λ = 561 nm, λ = 633 nm and λ = 405 nm lasers, respectively. Emitted signals were captured at λ = 569–614 nm, λ = 641–694 nm and λ = 410–520 nm for SBG-TMR, BG-SiR and Hoechst33342, respectively. Control experiments were performed in either SBG-TMR- or BG-SiR-labelled cells to exclude trafficking artefacts due to spectral overlap.

### Single molecule pulldown assay

Single molecule pulldown (SiMPull) was performed using HA-tagged GPCRs isolated on glass coverslips as previously described using a biotinylated anti-HA antibody^42^. Briefly, flow chambers were prepared with mPEG-passivated glass slides and coverslips with ∼1% biotinylated PEG to allow antibody capture. Prior to each experiment, flow chambers were incubated with 0.2 mg/ml NeutrAvidin for 2 min then incubated with 10 nM of antibody (abcam, ab26228) for 30 min. The flow chambers were rinsed with T50 buffer (50 mM NaCL, 10 mM Tris, pH 7.5) after each conjugation step. Cell lysate was prepared 24-48 hrs after transfection with HEK293T cells and immediately after labeling at 37 °C with either with 1 µM SBG-JF_549_ for 45 minutes or 20 µM SNAP-Surface® Block (NEB) followed by 1 µM BG-JF_549_ for 45 min each. After extensive washing with EX solution (in mM, 120 KCl, 29 NaCl, 1 MgCl_2_, 2 CaCl_2_ and 10 HEPES, pH 7.4), cells were harvested using Ca^2+^ free-DPBS for 20 min at 37 °C. After pelleting the cells at 10,000 x g, 4 °C for 1 min, cells were lysed using 1.2% IGEPAL detergent for 1 hour at 4 °C. Next, cells were centrifuged at 16,000 x g for 20 min at 4 °C and supernatant was collected and stored in ice until used. The cell lysate samples were then diluted using a dilution buffer containing 0.1% IGEPAL and introduced to the flow chamber. After obtaining an optimal number of spots in the field of view, the chamber was washed with the dilution buffer to remove unbound proteins.

Single molecule imaging was done using a 100x objective (NA 1.49) on an inverted microscope (Olympus IX83) in total internal reflection (TIR) mode at 20 Hz with 50 ms exposure times with an scMOS camera (Hamamatsu ORCA-Flash4v3.0). Samples were excited with 561 nm lasers and imaged using an emission filters of 595±25 nm. Data analysis was performed using custom made LabVIEW program as previously described^49^. Data was collected across at least 2 separate experimental days and then averaged to produce bar graphs in **Fig. 5d** and **5h**.

## Acknowledgements

We thank Andrea Bergner for providing purified SNAPf protein, Kai Johnsson and Birgit Koch for SNAP plasmids, Sebastian Fabritz and Cornelia Ullrich for small molecule and full protein mass spectrometry, Kai Johnsson for support and Philipp Leippe for helpful discussion (all MPIMR). D.J.H. was supported by a Diabetes UK R.D. Lawrence (12/0004431) Fellowship, a Wellcome Trust Institutional Support Award, MRC Confidence in Concept, and MRC (MR/N00275X/1 and MR/S025618/1) and Diabetes UK (17/0005681) Project Grants. This project has received funding from the European Research Council (ERC) under the European Union’s Horizon 2020 research and innovation programme (Starting Grant 715884 to D.J.H.). JL is supported by an R35 grant (1 R35 GM124731) from NIGMS and the Rohr Family Research Scholar Award.

## Author Contributions

DJH, JL and JB conceived and supervised the study. JB designed and PP, BM and JB synthesized and characterized all compounds. BJJ provided reagents. ED’E and JB performed STED nanoscopy. JB recorded *in vitro* solubility and labelling assays. VG, JA, DJH, JLee, JL and JB performed labelling and microscopy of GPCRs. JLee performed and analyzed SiMPull. VG performed and analyzed *in vivo* labelling. DJH, JL and JB wrote the manuscript with input from all authors.

