## Supplemental Information for "Interrogating surface versus intracellular transmembrane receptor populations using cell-impermeable SNAP-tag substrates"

**Table of Contents**

**1. General** 3

**2. Synthesis** 4

*2.1. Synthesis for CBG-NHCOCF$_3$* 4

*2.2. 6-((4-(Aminomethyl)benzyl)oxy)pyrimidine-2,4-diamine (3)* 4

*2.3. N-(4-(((2,6-Diaminopyrimidin-4-yl)oxy)methyl)benzyl)-2,2,2-trifluoroacetamide (4)* 4

*2.4. N-(4-(((2,6-Diamino-5-nitrosopyrimidin-4-yl)oxy)methyl)benzyl)-2,2,2-trifluoroacetamide (5)* 5

*2.5. Methyl 4-((2-amino-5-nitroso-6-((4-((2,2,2-trifluoroacetamido)methyl)benzyl)oxy)pyrimidin-4-yl)amino)-4-oxobutanoate (6)* 5

*2.6. Methyl 4-((2,5-diamino-6-((4-((2,2,2-trifluoroacetamido)methyl)benzyl)oxy)pyrimidin-4-yl)amino)-4-oxobutanoate (7)* 6

*2.7. Methyl 3-(2-amino-6-((4-((2,2,2-trifluoroacetamido)methyl)benzyl)oxy)-9H-purin-8-yl)propanoate (8)* 6

*2.8. 3-(2-Amino-6-((4-((2,2,2-trifluoroacetamido)methyl)benzyl)oxy)-9H-purin-8-yl)propanoic acid (CBG-NHCOCF$_3$)* 7

*2.9. 3-(2-Amino-6-((4-(aminomethyl)benzyl)oxy)-9H-purin-8-yl)propanoic acid (CBG-NH$_2$)* 7

*2.10. 2-(3-(2-Amino-6-((4-((2,2,2-trifluoroacetamido)methyl)benzyl)oxy)-9H-purin-8-yl)propanamido)ethane-1-sulfonic acid (SBG-NHCOCF$_3$)* 8

*2.11. 2-(3-(2-Amino-6-((4-(aminomethyl)benzyl)oxy)-9H-purin-8-yl)propanamido)ethane-1-sulfonic acid (SBG-NH$_2$)* 8

*2.12. General procedure A for NHS activated fluorophores* 8

*2.13. General procedure B for BG, CBG and SBG conjugates* 9

*2.14. 4-((4-(((2-Amino-9H-purin-6-yl)oxy)methyl)benzyl)carbamoyl)-2-(2,7-difluoro-6-hydroxy-3-oxo-3H-xanthen-9-yl)benzoate (BG-OG)* 9

*2.15. 4-((4-(((2-Amino-9H-purin-6-yl)oxy)methyl)benzyl)carbamoyl)-2-(6-(dimethylamino)-3-(dimethyliminio)-3H-xanthen-9-yl)benzoate (BG-TMR)* 9

*2.16. 4-((4-(((2-Amino-9H-purin-6-yl)oxy)methyl)benzyl)carbamoyl)-2-(3-(azetidin-1-ium-1-ylidene)-6-(azetidin-1-yl)-3H-xanthen-9-yl)benzoate (BG-JF$_{549}$)* 9

*2.17. 4-((4-(((2-Amino-9H-purin-6-yl)oxy)methyl)benzyl)carbamoyl)-2-(7-(dimethylamino)-3-(dimethyliminio)-5,5-dimethyl-3,5-dihydrodibenzo[b,e]silin-10-yl)benzoate (BG-SiR)* 10

*2.18. 4-((4-(((2-Amino-9H-purin-6-yl)oxy)methyl)benzyl)carbamoyl)-2-(3-(azetidin-1-ium-1-ylidene)-7-(azetidin-1-yl)-5,5-dimethyl-3,5-dihydrodibenzo[b,e]silin-10-yl)benzoate (BG-JF$_{646}$) 10*
*2.19. 4-((4-(((2-Amino-8-(2-carboxyethyl)-9H-purin-6-yl)oxy)methyl)benzyl)carbamoyl)-2-(2,7-difluoro-6-hydroxy-3-oxo-3H-xanthen-9-yl)benzoate (CBG-OG) 10*
*2.20. 4-((4-(((2-Amino-8-(2-carboxyethyl)-9H-purin-6-yl)oxy)methyl)benzyl)carbamoyl)-2-(6-(dimethylamino)-3-(dimethyliminio)-3H-xanthen-9-yl)benzoate (CBG-TMR) 11*
*2.21. 4-((4-(((2-Amino-8-(2-carboxyethyl)-9H-purin-6-yl)oxy)methyl)benzyl)carbamoyl)-2-(3-(azetidin-1-ium-1-ylidene)-6-(azetidin-1-yl)-3H-xanthen-9-yl)benzoate (CBG-JF$_{549}$) 11*
*2.22. 4-((4-(((2-Amino-8-(2-carboxyethyl)-9H-purin-6-yl)oxy)methyl)benzyl)carbamoyl)-2-(7-(dimethylamino)-3-(dimethyliminio)-5,5-dimethyl-3,5-dihydrodibenzo[b,e]silin-10-yl)benzoate (CBG-SiR) 11*
*2.23. 4-((4-(((2-Amino-8-(2-carboxyethyl)-9H-purin-6-yl)oxy)methyl)benzyl)carbamoyl)-2-(3-(azetidin-1-ium-1-ylidene)-7-(azetidin-1-yl)-5,5-dimethyl-3,5-dihydrodibenzo[b,e]silin-10-yl)benzoate (CBG-JF$_{646}$) 12*
*2.24. 4-((4-(((2-Amino-8-(3-oxo-3-((2-sulfoethyl)amino)propyl)-9H-purin-6-yl)oxy)methyl)benzyl)carbamoyl)-2-(2,7-difluoro-6-hydroxy-3-oxo-3H-xanthen-9-yl)benzoate (SBG-OG) 12*
*2.25. 4-((4-(((2-Amino-8-(3-oxo-3-((2-sulfoethyl)amino)propyl)-9H-purin-6-yl)oxy)methyl)benzyl)carbamoyl)-2-(6-(dimethylamino)-3-(dimethyliminio)-3H-xanthen-9-yl)benzoate (SBG-TMR) 12*
*2.26. 4-((4-(((2-Amino-8-(3-oxo-3-(2-sulfoethoxy)propyl)-9H-purin-6-yl)oxy)methyl)benzyl)carbamoyl)-2-(3-(azetidin-1-ium-1-ylidene)-6-(azetidin-1-yl)-3H-xanthen-9-yl)benzoate (SBG-JF$_{549}$) 13*
*2.27. 4-((4-(((2-Amino-8-(3-oxo-3-(2-sulfoethoxy)propyl)-9H-purin-6-yl)oxy)methyl)benzyl)carbamoyl)-2-(7-(dimethylamino)-3-(dimethyliminio)-5,5-dimethyl-3,5-dihydrodibenzo[b,e]silin-10-yl)benzoate (SBG-SiR) 13*
*2.28. 4-((4-(((2-Amino-8-(3-oxo-3-((2-sulfoethyl)amino)propyl)-9H-purin-6-yl)oxy)methyl)benzyl)carbamoyl)-2-(3-(azetidin-1-ium-1-ylidene)-7-(azetidin-1-yl)-5,5-dimethyl-3,5-dihydrodibenzo[b,e]silin-10-yl)benzoate (SBG-JF$_{646}$) 13*

**3. SNAP$_f$ construct and mass spectrometry 14**

**4. Supplementary Schemes 15**

**5. Supplementary Figures: 17**

**6. References 22**

### 1. General

All chemical reagents and anhydrous solvents for synthesis were purchased from commercial suppliers (Sigma-Aldrich, Fluka, Acros, Fluorochem, TCI) and were used without further purification or distillation. If necessary, solvents were degassed either by freeze-pump-thaw or by bubbling $N_2$ through the vigorously stirred solution for several minutes.

NMR spectra were recorded in deuterated solvents on a Bruker AVANCE III HD 400 equipped with a CryoProbe and calibrated to residual solvent peaks ($^1H/^{13}C$ in ppm): $CDCl_3$ (7.26/77.00), DMSO-$d_6$ (2.50/39.52), acetone-$d_6$ (2.05/29.84), MeOD-$d_4$ (3.31/49.00), DMF-$d_7$ (8.03/163.15. Multiplicities are abbreviated as follows: s = singlet, d = doublet, t = triplet, q = quartet, p = pentet, br = broad, m = multiplet. Coupling constants *J* are reported in Hz. Spectra are reported based on appearance, not on theoretical multiplicities derived from structural information.

LC-MS was performed on a Shimadzu MS2020 connected to a Nexera UHPLC system equipped with a Waters ACQUITY UPLC BEH C18 (1.7 µm, 50 × 2.1 mm). Buffer A: 0.1% FA in H2O Buffer B: acetonitrile. The typical gradient was from 10% B for 0.5 min → gradient to 90% B over 4.5 min → 90% B for 0.5 min → gradient to 99% B over 0.5 min with 1 mL/min flow. Retention times (tR) are given in minutes (min).

High resolution mass spectrometry was performed using a Bruker maXis II ETD hyphenated with a Shimadzu Nexera system. The instruments were controlled via Brukers otofControl 4.1 and Hystar 4.1 SR2 (4.1.31.1) software. The acquisition rate was set to 3 Hz and the following source parameters were used for positive mode electrospray ionization: End plate offset = 500 V; capillary voltage = 3800 V; nebulizer gas pressure = 45 psi; dry gas flow = 10 L/min; dry temperature = 250 °C. Transfer, quadrupole and collision cell settings are mass range dependent and were fine-adjusted with consideration of the respective analyte's molecular weight. For internal calibration sodium format clusters were used. Samples were desalted via fast liquid chromatography. A Supelco TitanTM C18 UHPLC Column, 1.9 µm, 80 Å pore size, 20 × 2.1 mm and a 2 min gradient from 10 to 98% aqueous MeCN with 0.1% FA ($H_2O$: Carl Roth GmbH + Co. KG ROTISOLV® Ultra LC-MS; MeCN: Merck KGaA LiChrosolv® Acetonitrile hypergrade for LC-MS; FA - Merck KGaA LiChropur® Formic acid 98%- 100% for LC-MS) was used for separation. Sample dilution in 10% aqueous ACN (hyper grade) and injection volumes were chosen dependent of the analyte's ionization efficiency. Hence, on-column loadings resulted betwee 0.25–5.0 ng. Automated internal re-calibration and data analysis of the recorded spectra were performed with Bruker's DataAnalysis 4.4 SR1 software.

Preparative RP-HPLC was performed on a Waters e2695 system equipped with a 2998 PDA detector for product collection (at 220, or 650 nm) on a Supelco Ascentis® C18 HPLC Column (5 µm, 250 × 21.2 mm). Buffer A: 0.1% TFA in $H_2O$ Buffer B: MeCN. The typical gradient was from 10% B for 5 min → gradient to 90% B over 45 min → 90% B for 5 min → gradient to 99% B over 5 min with 8 mL/min flow.

Flash column chromatography was performed on a Biotage Isolera One with pre-packed silica columns (0.040–0.063 mm, 230-400 mesh, Silicycle). Reactions and chromatography fractions were monitored by thin layer chromatography (TLC) on Merck silica gel 60 F254 glass plates. The spots were visualized either under UV light at 254 nm and/or 366 nm or with appropriate staining method (iodine, *para*-anisaldehyde, $KMnO_4$) followed by heating.

### 2. Synthesis

#### 2.1. Synthesis for CBG-$NHCOCF_3$

CBG-$NHCOCF_3$ was prepared as described previously in 7 steps[1] from 6-chloropyrimidine-2,4-diamine and (4-(aminomethyl)phenyl)methanol in a slightly modified procedure. Each step was characterized by LCMS and LRMS is reported below.

#### 2.2. 6-((4-(Aminomethyl)benzyl)oxy)pyrimidine-2,4-diamine (3)

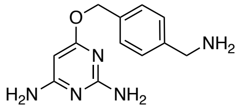


A flame-dried Schlenk flask was charged with 2.00 g (14.59 mmol, 1.0 equiv.) of 4-(aminomethyl)-benzylalcohol (**2**) under a nitrogen atmosphere and dissolved in 25 mL of dry pyridine and 25 mL of dry xylene. 1.16 g (29.00 mmol, 2.0 equiv.) of NaH (60 % in mineral oil) were added in portions under a stream of nitrogen. The reaction mixture was heated to 120 °C until all NaH was dissolved, upon which 2.25 g (15.61 mmol, 1.07 equiv.) of 2,4-diamino-6-chloropyrimidin (**1**) were added slowly. A reflux condenser was attached and the brown suspension was stirred for 3 h at 150 °C under a nitrogen atmosphere before it was cooled to r.t. and filtered over a por.4 glass frit to remove remaining solids. The solid was rinsed with 20 mL of EtOAc and the combined filtrate was dried *in vacuo* and directly used in next step without further purification.

**LRMS** (ESI): calc. for $C_{13}H_{16}N_5O$ $[M+H]^+$: 246.1, found: 246.0.

#### 2.3. *N*-(4-(((2,6-Diaminopyrimidin-4-yl)oxy)methyl)benzyl)-2,2,2-trifluoroacetamide (4)

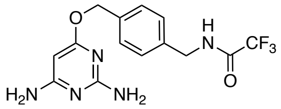


Crude **3** (theoretical 14.59 mmol) was dissolved in 40 mL of ethanol and 2 mL of DIPEA. 9.8 mL (72.95 mmol) of trifluoroethyl-trifluoracetat were added at r.t. and the reaction mixture was stirred for 16 h before it was evaporated to total dryness, taken up in 10% MeOH in DCM and filtered over a plug of silica with extensive rinsing to afford 2.28 g (6.69 mmol) of the desired product as a brownish solid in 46% yield over 2 steps.

**LRMS** (ESI): calc. for $C_{14}H_{15}F_3N_5O_2$ $[M+H]^+$: 342.3, found: 342.0.

### 2.4. *N*-(4-(((2,6-Diamino-5-nitrosopyrimidin-4-yl)oxy)methyl)benzyl)-2,2,2-trifluoroacetamide (5)

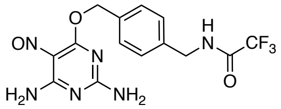


In a round bottom flask, 1.58 g (4.63 mmol) of **4** were dissolved in 10 mL of acetic acid and 22 mL of water. The yellowish solution was heated to 70 °C before 830 mg (12.0 mmol, 2.6 equiv.) of $NaNO_2$ in 10 mL of water were slowly added dropwise. The reaction mixture turns into a deep purple suspension that was stirred for 1 h at 70 °C before the reaction mixture was cooled to rt and stirred for another 30 min. The purple solid was filtered and rinsed with 20 mL of cold water, suspended in 20 mL MeCN:water = 1:1 and lyophilized to obtain 966 mg (2.60 mmol) of the desired product as a purple solid in 56% yield.

**$^1$H NMR** (400 MHz, DMSO-d$_6$): $\delta$ [ppm] = 10.41–9.76 (m, 2H), 8.02 (d, $J$ = 4.7 Hz, 2H), 7.84 (m, 2H), 7.52 (d, $J$ = 7.9 Hz, 2H), 7.31 (d, $J$ = 8.1 Hz, 2H), 5.56 (s, 2H), 4.40 (d, $J$ = 5.9 Hz, 2H).
**LRMS** (ESI): calc. for $C_{14}H_{14}F_3N_6O_3$ $[M+H]^+$: 371.1, found: 370.9.

### 2.5. Methyl 4-((2-amino-5-nitroso-6-((4-((2,2,2-trifluoroacetamido)methyl)benzyl)oxy)pyrimidin-4-yl)amino)-4-oxobutanoate (6)

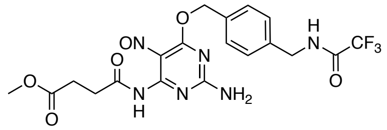


In a round bottom flask, 1.20 g (3.24 mmol, 1.0 equiv.) of **5** was dissolved in 20 mL of dry acetone and 1.13 mL (6.48 mmol, 2.0 equiv.) of DIPEA before 440 µL (3.56 mmol, 1.1 equiv.) of methyl-4-chloro-4-oxobutyrate was added dropwise to solution. The initially blue reaction mixture was stirred for 1 h at room temperature until the solution turned green. 20 mL of MeOH was added, and the solvents were removed *in vacuo*. The green solid was purified by filtration over silica gel with 10% MeOH in DCM.

**LRMS** (ESI): calc. for $C_{19}H_{20}F_3N_6O_6$ $[M+H]^+$: 485.1, found: 485.1.

### 2.6. Methyl 4-((2,5-diamino-6-((4-((2,2,2-trifluoroacetamido)methyl)benzyl)oxy)pyrimidin-4-yl)amino)-4-oxobutanoate (7)

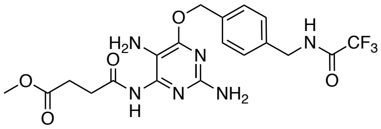


In a Schlenk flask under an argon atmosphere, crude **6** was dissolved in 20 mL of degassed ethanol and a catalytic amount of Raney nickel (suspension in water) was added. The mixture was purged with 3 ballons of $H_2$ and stirred for 1 h at r.t. until the color changed to yellow. Raney nickel was removed by filtration using a syringe filter (0.45 µm) and the filtrate was used in the next step without further purification.

**LRMS** (ESI): calc. for $C_{19}H_{22}F_3N_6O_5$ $[M+H]^+$: 471.2, found: 471.0.

### 2.7. Methyl 3-(2-amino-6-((4-((2,2,2-trifluoroacetamido)methyl)benzyl)oxy)-9*H*-purin-8-yl)propanoate (8)

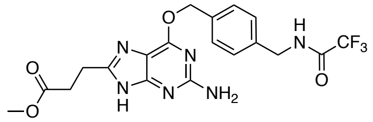


In a round bottom flask, 647 µL of acetic acid were added to crude **7** and the solution was refluxed for 2 h. 20 mL of toluene was added and the solvents were removed *in vacuo*. The remaining yellowish solid was taken up in DMF/water = 8/2 and purified by RP-HPLC (MeCN:water + 0.1% TFA, 10-90 % MeCN over 60 min, flow 8 mL/min, λ = 293 nm) to obtain 360 mg (0.80 mmol) of desired product as a yellowish solid in 25% yield over 3 steps.

**$^1$H NMR** (400 MHz, DMSO-d$_6$): $\delta$ [ppm] = 12.20 (s, 1H), 10.00 (t, $J$ = 6.0 Hz, 1H), 7.48 (d, $J$ = 7.8 Hz, 2H), 7.30 (d, $J$ = 7.9 Hz, 2H), 6.20 (s, 2H), 5.44 (s, 2H), 4.40 (d, $J$ = 5.9 Hz, 2H), 3.58 (s, 3H), 2.92 (t, $J$ = 7.3 Hz, 2H), 2.79 (t, $J$ = 7.2 Hz, 2H).
**HRMS** (ESI): calc. for $C_{19}H_{20}F_3N_6O_4$ $[M+H]^+$: 453.1493, found: 453.1495.

#### 2.8. 3-(2-Amino-6-((4-((2,2,2-trifluoroacetamido)methyl)benzyl)oxy)-9*H*-purin-8-yl)propanoic acid (CBG-NHCOCF$_3$)

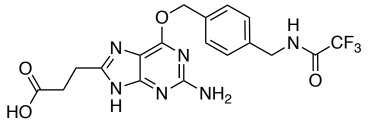


A 40 mL dram vial was charged with 360 mg (0.80 mmol, 1.0 equiv.) of **8** dissolved in 10 mL of MeOH with the addition of 0.1 mL 1.5 M NaOH and cooled to 0 °C. Once cooled, 2.20 g (39.8 mmol, 50.0 equiv.) of KOH was added in one portion and the reaction mixture was stirred vigorously for 5 h while maintained at 0 °C before it was quenched by addition 3.5 mL HOAc. All volatiles were removed *in vacuo* and the residue was taken up in DMF:water:MeCN (7 mL total, 5:1:1) and purified by RP-HPLC (MeCN:water + 0.1% TFA, 10-90 % MeCN over 60 min, flow 8 mL/min, $\lambda$ = 293 nm) to obtain 190 mg (0.43 mmol) of the desired product as a white solid in 54% yield, plus 20 mg (0.06 mmol, 7%) of **CBG-NH$_2$** and 25 mg (0.06 mmol, 7%) of recovered starting material.

**$^1$H NMR** (400 MHz, DMF-d$_7$): $\delta$ [ppm] = 10.03 (t, $J$ = 6.0 Hz, 1H), 7.59 (d, $J$ = 7.9 Hz, 2H), 7.42 (d, $J$ = 7.9 Hz, 2H), 5.56 (s, 2H), 4.55 (d, $J$ = 6.1 Hz, 2H), 3.15 (t, $J$ = 7.2 Hz, 2H), 2.90 (t, $J$ = 6.6 Hz, 2H).
**HRMS** (ESI): calc. for $C_{18}H_{28}F_3N_6O_4$ $[M+H]^+$: 439.1336, found: 439.1335.

#### 2.9. 3-(2-Amino-6-((4-(aminomethyl)benzyl)oxy)-9*H*-purin-8-yl)propanoic acid (CBG-NH$_2$)

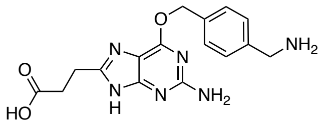


**CBG-NH$_2$** was isolated alongside **CBG-NHCOCF$_3$** in 7% yield.

**$^1$H NMR** (400 MHz, DMSO-d$_6$): $\delta$ [ppm] = 10.03 (t, $J$ = 6.1 Hz, 1H), 8.25 (s, 2H), 7.68–7.43 (m, 2H), 7.43–7.18 (m, 2H), 5.49 (s, 2H), 4.40 (d, $J$ = 6.0 Hz, 2H), 2.98 (t, $J$ = 7.2 Hz, 2H), 2.75 (t, $J$ = 7.2 Hz, 2H), 2.55 (t, $J$ = 5.5 Hz, 4H).
**HRMS** (ESI): calc. for $C_{16}H_{19}N_6O_3$ $[M+H]^+$: 343.1513, found: 343.1511.

### 2.10. 2-(3-(2-Amino-6-((4-((2,2,2-trifluoroacetamido)methyl)benzyl)oxy)-9*H*-purin-8-yl)propanamido)ethane-1-sulfonic acid (SBG-NHCOCF₃)

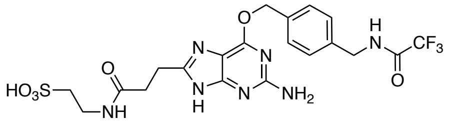

In a 4 mL dram vial, 13 mg (30 µmol, 1.0 equiv.) of **CBG-NHCOCF₃** was dissolved in 1.6 mL DMF and 22.5 µL DIPEA and stirred for 10 min. 30 mg (0.24 mmol, 8.0 equiv.) of taurine and 11.8 mg (39 µmol, 1.3 equiv.) of TSTU were added in one portion and the resulting suspension was stirred for 2 h at r.t.. 400 µL of water were added and the mixture was purified by RP-HPLC (MeCN:water + 0.1% TFA, 10-90 % MeCN over 60 min, flow 8 mL/min, λ = 293 nm) to obtain 10 mg (18 µmol) of the desired product as a white powder after freeze-drying in 61% yield.

**HRMS** (ESI): calc. for $C_{20}H_{23}F_3N_7O_6S$ $[M+H]^+$: 546.1377, found: 546.1374.

### 2.11. 2-(3-(2-Amino-6-((4-(aminomethyl)benzyl)oxy)-9*H*-purin-8-yl)propanamido)ethane-1-sulfonic acid (SBG-NH₂)

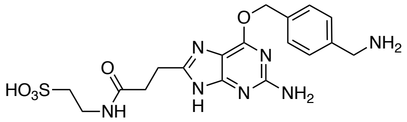

In a 40 mL dram vial, 10 mg (0.018 mmol) of **SBG-NHCOCF₃** was dissolved in 11.5 mL MeOH and 765 µL water before 245 mg (1.77 mmol) of $K_2CO_3$ was added in one portion. The suspension was stirred at 80 °C for 1 h before it was concentrated to a volume of ~300 µL *in vacuo*. 300 µL DMF, 300 µL MeOH and 300 µL water were added and the mixture was purified by RP-HPLC (MeCN:water + 0.1% TFA, 1-40% MeCN over 60 min, flow 8 mL/min, λ = 293 nm) to obtain 8.5 mg (0.015 mmol) of the desired product as a TFA salt after freeze-drying in 87% yield.

**$^1$H NMR** (400 MHz, DMSO-d₆): $\delta$ [ppm] = 8.13 (br s, 3H), 7.86 (t, $J$ = 5.5 Hz, 1H), 7.58 (d, $J$ = 8.0 Hz, 2H), 7.48 (d, $J$ = 8.0 Hz, 2H), 5.55 (s, 2H), 4.17–3.92 (m, 2H), 3.36–3.13 (m, 2H), 2.99 (t, $J$ = 7.1 Hz, 2H), 2.59 (t, $J$ = 7.1 Hz, 2H), 2.52 (t, $J$ = 7.5 Hz, 2H).
**HRMS** (ESI): calc. for $C_{18}H_{24}N_7O_5S$ $[M+H]^+$: 450.1554, found: 450.1556.

### 2.12. General procedure A for NHS activated fluorophores

In an Eppendorf tube, 3.5 µmol of carboxy-dye was dissolved in 100 µL DMF and 5 µL of DIPEA. Upon addition of 116 µL of TSTU (10 mg/mL in DMF, 3.86 µmol) the reaction mixture was vortexed and allowed to incubate for 10 min, before 40 µL of acetic acid and 200 µL of water were added and the solution was subjected to RP-HPLC (MeCN:water + 0.1% TFA, 10-90% MeCN over 60 min, flow 8 mL/min, threshold = 20). The product containing fractions were pooled and lyophilized.

### 2.13. General procedure B for BG, CBG and SBG conjugates

In an Eppendorf tube, 3.5 µmol of NHS-activated-dye from general procedure A was dissolved in 100 µL DMF and 5 µL of DIPEA, to which 3.5 µmol of amine (*i.e.* BG, CBG or SBG) dissolved in 200 µL DMF:water mixture (7:3) was added and the mixture was stirred for 2 h, before 40 µL of acetic acid and 200 µL of water were added and the solution was subjected to RP-HPLC (MeCN:water + 0.1% TFA, 10-90% MeCN over 60 min, flow 8 mL/min, threshold = 20). The product containing fractions were pooled and lyophilized to obtain the desired product as a colourful powder.

### 2.14. 4-((4-(((2-Amino-9*H*-purin-6-yl)oxy)methyl)benzyl)carbamoyl)-2-(2,7-difluoro-6-hydroxy-3-oxo-3*H*-xanthen-9-yl)benzoate (BG-OG)

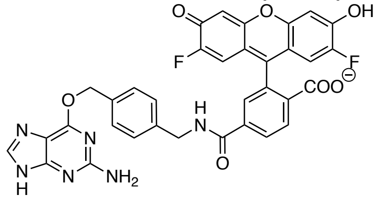

**BG-OG** was prepared according to general procedure A and B.
**HPLC**: λ = 500 nm.
**HRMS** (ESI): calc. for $C_{34}H_{24}F_2N_6O_7$ $[M+2H]^{2+}$: 333.0832, found: 333.0833.

### 2.15. 4-((4-(((2-Amino-9*H*-purin-6-yl)oxy)methyl)benzyl)carbamoyl)-2-(6-(dimethylamino)-3-(dimethyliminio)-3*H*-xanthen-9-yl)benzoate (BG-TMR)

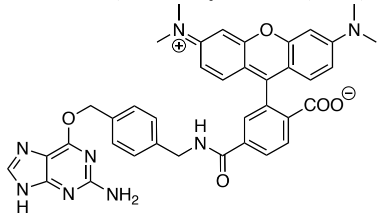

**BG-TMR** was prepared according to general procedure A and B.
**HPLC**: λ = 550 nm.
**HRMS** (ESI): calc. for $C_{38}H_{36}N_8O_5$ $[M+2H]^{2+}$: 342.1399, found: 342.1399.

### 2.16. 4-((4-(((2-Amino-9*H*-purin-6-yl)oxy)methyl)benzyl)carbamoyl)-2-(3-(azetidin-1-ium-1-ylidene)-6-(azetidin-1-yl)-3*H*-xanthen-9-yl)benzoate (BG-JF$_{549}$)

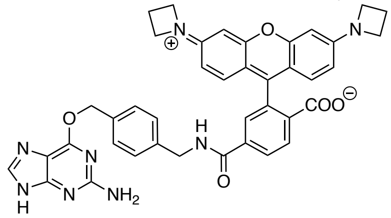

**BG-JF$_{549}$** was prepared according to general procedure A and B.
**HPLC**: λ = 550 nm.
**HRMS** (ESI): calc. for $C_{40}H_{36}N_8O_5$ $[M+2H]^{2+}$: 354.1399, found: 354.1397.

#### 2.17. 4-((4-(((2-Amino-9*H*-purin-6-yl)oxy)methyl)benzyl)carbamoyl)-2-(7-(dimethylamino)-3-(dimethyliminio)-5,5-dimethyl-3,5-dihydrodibenzo[*b*,*e*]silin-10-yl)benzoate (BG-SiR)

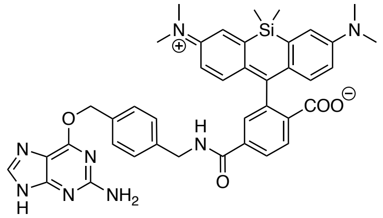

**BG-SiR** was prepared according to general procedure A and B.
**HPLC**: λ = 650 nm.
**HRMS** (ESI): calc. for $C_{40}H_{42}N_8O_4Si$ $[M+2H]^{2+}$: 363.1544, found: 363.1544.

#### 2.18. 4-((4-(((2-Amino-9*H*-purin-6-yl)oxy)methyl)benzyl)carbamoyl)-2-(3-(azetidin-1-ium-1-ylidene)-7-(azetidin-1-yl)-5,5-dimethyl-3,5-dihydrodibenzo[*b*,*e*]silin-10-yl)benzoate (BG-JF$_{646}$)

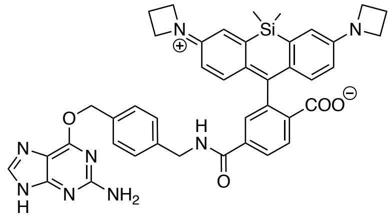

**BG- JF$_{646}$** was prepared according to general procedure A and B.
**HPLC**: λ = 650 nm.
**HRMS** (ESI): calc. for $C_{42}H_{42}N_8O_4Si$ $[M+2H]^{2+}$: 375.1544, found: 375.1543.

#### 2.19. 4-((4-(((2-Amino-8-(2-carboxyethyl)-9*H*-purin-6-yl)oxy)methyl)benzyl)carbamoyl)-2-(2,7-difluoro-6-hydroxy-3-oxo-3*H*-xanthen-9-yl)benzoate (CBG-OG)

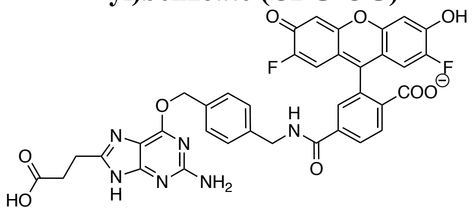

**CBG-OG** was prepared according to general procedure A and B.
**HPLC**: λ = 500 nm.
**HRMS** (ESI): calc. for $C_{37}H_{28}F_2N_6O_9$ $[M+2H]^{2+}$: 369.0937, found: 369.0939.

**2.20. 4-((4-(((2-Amino-8-(2-carboxyethyl)-9*H*-purin-6-yl)oxy)methyl)benzyl)carbamoyl)-2-(6-(dimethylamino)-3-(dimethyliminio)-3*H*-xanthen-9-yl)benzoate (CBG-TMR)**

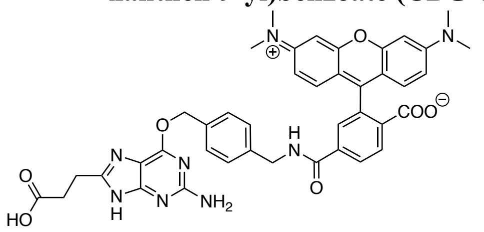

**CBG-TMR** was prepared according to general procedure A and B.
**HPLC**: λ = 550 nm.
**HRMS** (ESI): calc. for $C_{41}H_{40}N_8O_7$ $[M+2H]^{2+}$: 378.1504, found: 378.1507.

**2.21. 4-((4-(((2-Amino-8-(2-carboxyethyl)-9*H*-purin-6-yl)oxy)methyl)benzyl)carbamoyl)-2-(3-(azetidin-1-ium-1-ylidene)-6-(azetidin-1-yl)-3*H*-xanthen-9-yl)benzoate (CBG-JF$_{549}$)**

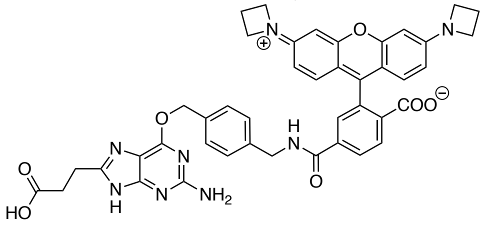

**CBG-JF$_{549}$** was prepared according to general procedure A and B.
**HPLC**: λ = 550 nm.
**HRMS** (ESI): calc. for $C_{43}H_{39}N_8O_7$ $[M+H]^{+}$: 779.2936, found: 779.2932.

**2.22. 4-((4-(((2-Amino-8-(2-carboxyethyl)-9*H*-purin-6-yl)oxy)methyl)benzyl)carbamoyl)-2-(7-(dimethylamino)-3-(dimethyliminio)-5,5-dimethyl-3,5-dihydrodibenzo[*b*,*e*]silin-10-yl)benzoate (CBG-SiR)**

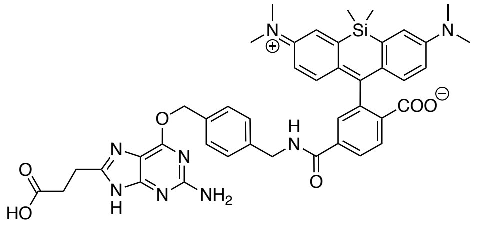

**CBG-SiR** was prepared according to general procedure A and B.
**HPLC**: λ = 650 nm.
**HRMS** (ESI): calc. for $C_{43}H_{46}N_8O_6Si$ $[M+2H]^{2+}$: 399.1649, found: 399.1649.

**2.23. 4-((4-(((2-Amino-8-(2-carboxyethyl)-9*H*-purin-6-yl)oxy)methyl)benzyl)carbamoyl)-2-(3-(azetidin-1-ium-1-ylidene)-7-(azetidin-1-yl)-5,5-dimethyl-3,5-dihydrodibenzo[*b*,*e*]silin-10-yl)benzoate (CBG-JF$_{646}$)**

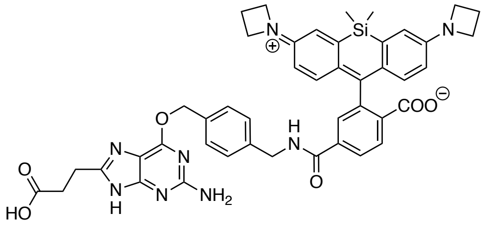

**CBG- JF$_{646}$** was prepared according to general procedure A and B.
**HPLC**: $\lambda$ = 650 nm.
**HRMS** (ESI): calc. for $C_{45}H_{46}N_8O_6Si$ $[M+2H]^{2+}$: 411.1649, found: 411.1646.

**2.24. 4-((4-(((2-Amino-8-(3-oxo-3-((2-sulfoethyl)amino)propyl)-9*H*-purin-6-yl)oxy)methyl)benzyl)carbamoyl)-2-(2,7-difluoro-6-hydroxy-3-oxo-3*H*-xanthen-9-yl)benzoate (SBG-OG)**

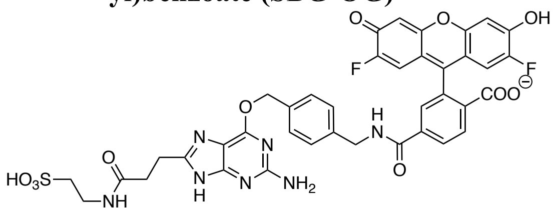

**SBG-OG** was prepared according to general procedure A and B.
**HPLC**: $\lambda$ = 500 nm.
**HRMS** (ESI): calc. for $C_{39}H_{32}F_2N_7O_{11}S$ $[M+H]^+$: 844.1843, found: 844.1843.

**2.25. 4-((4-(((2-Amino-8-(3-oxo-3-((2-sulfoethyl)amino)propyl)-9*H*-purin-6-yl)oxy)methyl)benzyl)carbamoyl)-2-(6-(dimethylamino)-3-(dimethyliminio)-3*H*-xanthen-9-yl)benzoate (SBG-TMR)**

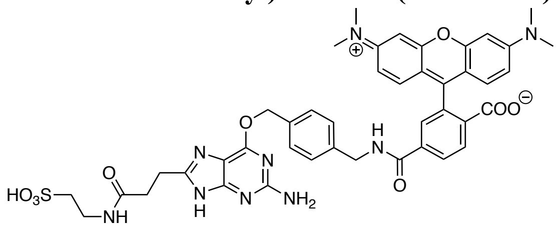

**SBG-TMR** was prepared according to general procedure A and B.
**HPLC**: $\lambda$ = 550 nm.
**HRMS** (ESI): calc. for $C_{43}H_{45}N_9O_9S$ $[M+2H]^{2+}$: 431.6525, found: 431.6519.

**2.26. 4-((4-(((2-Amino-8-(3-oxo-3-(2-sulfoethoxy)propyl)-9*H*-purin-6-yl)oxy)methyl)benzyl)carbamoyl)-2-(3-(azetidin-1-ium-1-ylidene)-6-(azetidin-1-yl)-3*H*-xanthen-9-yl)benzoate (SBG-JF$_{549}$)**

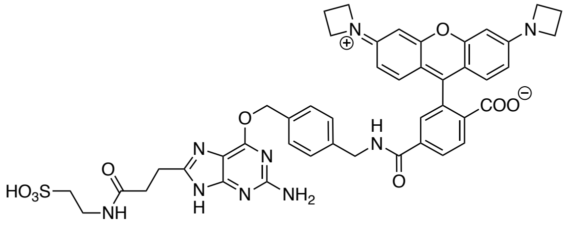

**SBG-JF$_{549}$** was prepared according to general procedure A and B.
**HPLC**: λ = 550 nm.
**HRMS** (ESI): calc. for $C_{45}H_{45}N_9O_9S$ $[M+2H]^{2+}$: 443.6525, found: 443.6522.

**2.27. 4-((4-(((2-Amino-8-(3-oxo-3-(2-sulfoethoxy)propyl)-9*H*-purin-6-yl)oxy)methyl)benzyl)carbamoyl)-2-(7-(dimethylamino)-3-(dimethyliminio)-5,5-dimethyl-3,5-dihydrodibenzo[*b*,*e*]silin-10-yl)benzoate (SBG-SiR)**

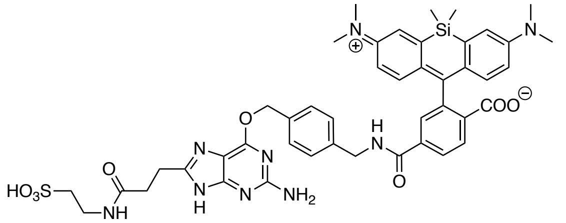

**SBG-SiR** was prepared according to general procedure A and B.
**HPLC**: λ = 650 nm.
**HRMS** (ESI): calc. for $C_{45}H_{51}N_9O_8SSi$ $[M+2H]^{2+}$: 452.6670, found: 452.6665.

**2.28. 4-((4-(((2-Amino-8-(3-oxo-3-((2-sulfoethyl)amino)propyl)-9*H*-purin-6-yl)oxy)methyl)benzyl)carbamoyl)-2-(3-(azetidin-1-ium-1-ylidene)-7-(azetidin-1-yl)-5,5-dimethyl-3,5-dihydrodibenzo[*b*,*e*]silin-10-yl)benzoate (SBG-JF$_{646}$)**

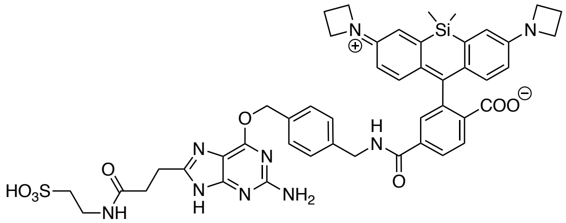

**SBG-JF$_{646}$** was prepared according to general procedure A and B.
**HPLC**: λ = 650 nm.
HRMS (ESI): calc. for $C_{47}H_{51}N_9O_8SSi$ $[M+2H]^{2+}$: 464.6670, found: 464.6667.

### 3. SNAP$_f$ construct and mass spectrometry

SNAP$_f$ sequence:

```
MASWSHPQFEKGADDDDKVPHMDKDCEMKRTTLDSPLGKLELSGCEQGLHRIIFLGKGTSAADAV
EVPAPAAVLGGPEPLMQATAWLNAYFHQPEAIEEFPVPALHHPVFQQESFTRQVLWKLLKVVKFG
EVISYSHLAALAGNPAATAAVKTALSGNPVPILIPCHRVVQGDLDVGGYEGGLAVKEWLLAHEGH
RLGKPGLGAPGFSSISAHHHHHHHHHH
```

Strep-Tag II, Enterokinase-site, SNAP$_f$, His-Tag

After His-tag purification, two posttranslational SNAP$_f$ constructs were observed: one with removed start codon SNAP$_f$ (2–222) and one without the N-terminal Strep-Tag II and Enterokinase-site SNAP$_f$ (22–222):

| Condition | calc. | found | Δ ppm |
|---|---|---|---|
| SNAP$_f$ (2–222) | 23865.101 | 23865.134 | 1.38 |
| SNAP$_f$ (22–222) | 21618.103 | 21618.132 | 1.34 |
| SNAP$_f$ (2–222):BG-TMR | 24396.317 | 24396.342 | 1.04 |
| SNAP$_f$ (22–222):BG-TMR | 22149.319 | 22149.347 | 1.26 |
| SNAP$_f$ (2–222):CBG-TMR | 24396.317 | 24396.333 | 0.66 |
| SNAP$_f$ (22–222):CBG-TMR | 22149.319 | 22149.336 | 0.79 |
| SNAP$_f$ (2–222):SBG-TMR | 24396.317 | 24396.338 | 0.86 |
| SNAP$_f$ (22–222):SBG-TMR | 22149.319 | 22149.344 | 1.14 |

### 4. Supplementary Schemes

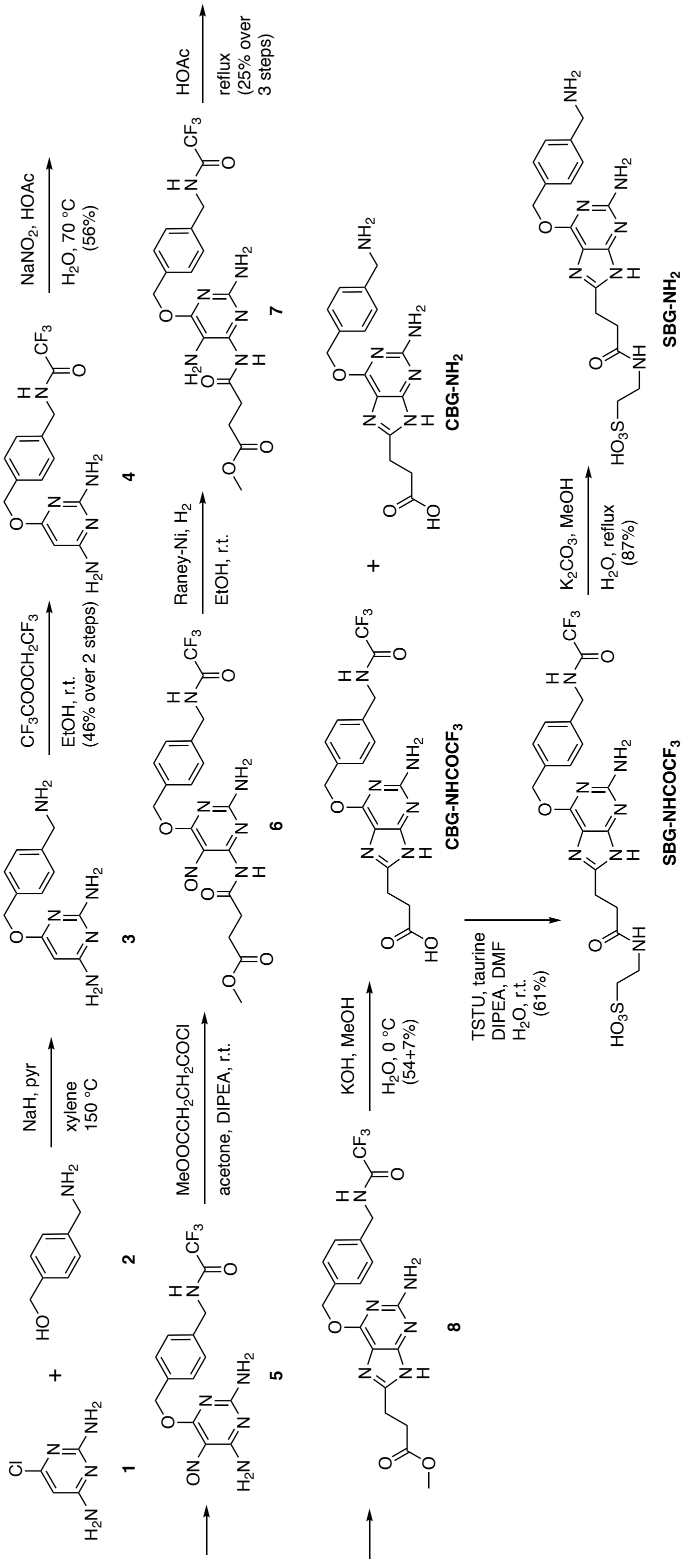


**Supplementary Scheme 1. Synthesis of SBG-NH$_2$.** CBG-NHCOCF$_3$ has been reported before[1], and was converted to SBG-NH$_2$ *via* peptide coupling to taurine and subsequent deprotection.

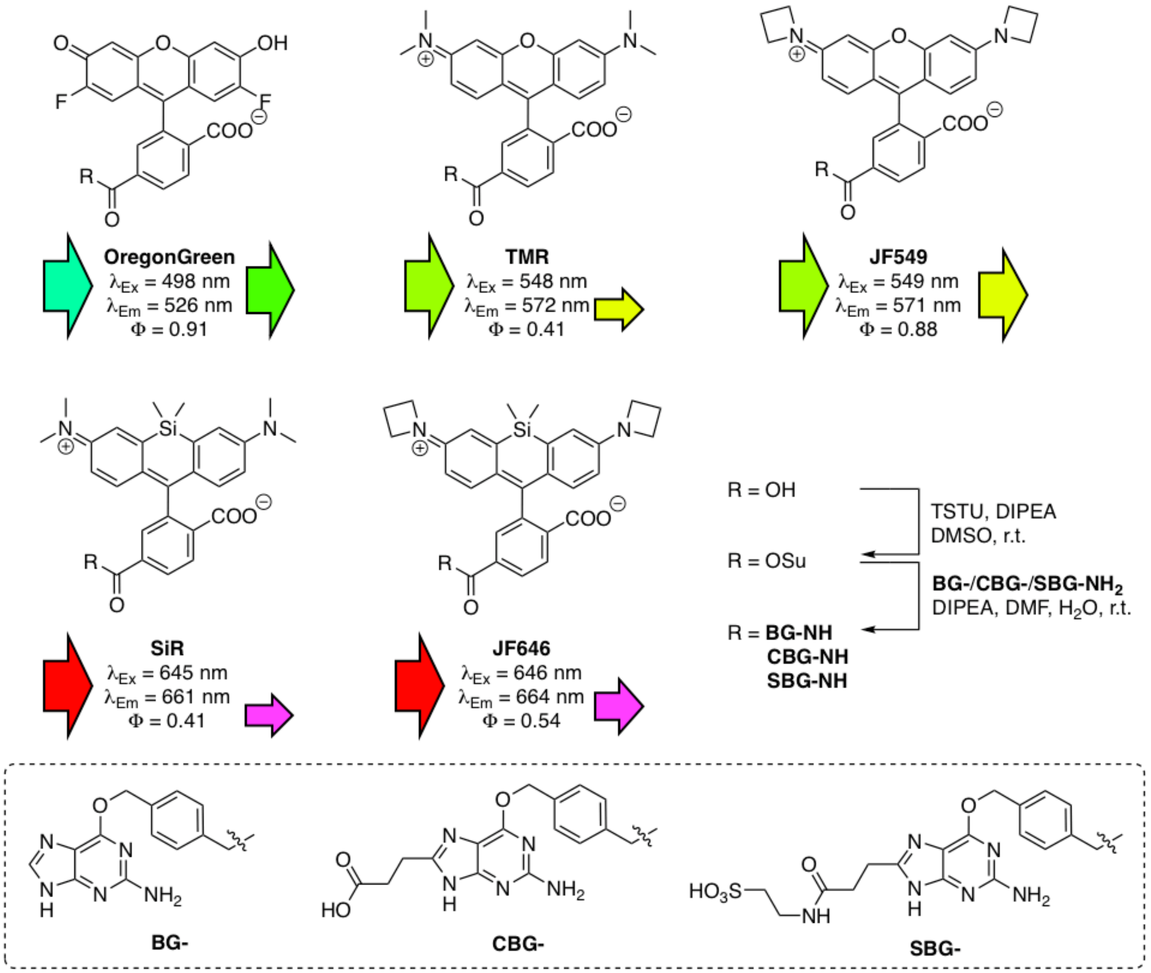


**Supplementary Scheme 2. Synthesis and properties of SBG-linked fluorophores.** Acid-containing fluorophores were activated with TSTU and the corresponding NHS ester isolated before reaction with SBG-NH2 to yield the desired impermeable dye. $\lambda_{Ex}$ = maximal absorption wavelength; $\lambda_{Em}$ = maximal emission wavelength; $\Phi$ = quantum yield. Left arrow represents color for excitation, right arrow represents color for emission, with the respective size depicting quantum yield.

### 5. Supplementary Figures:

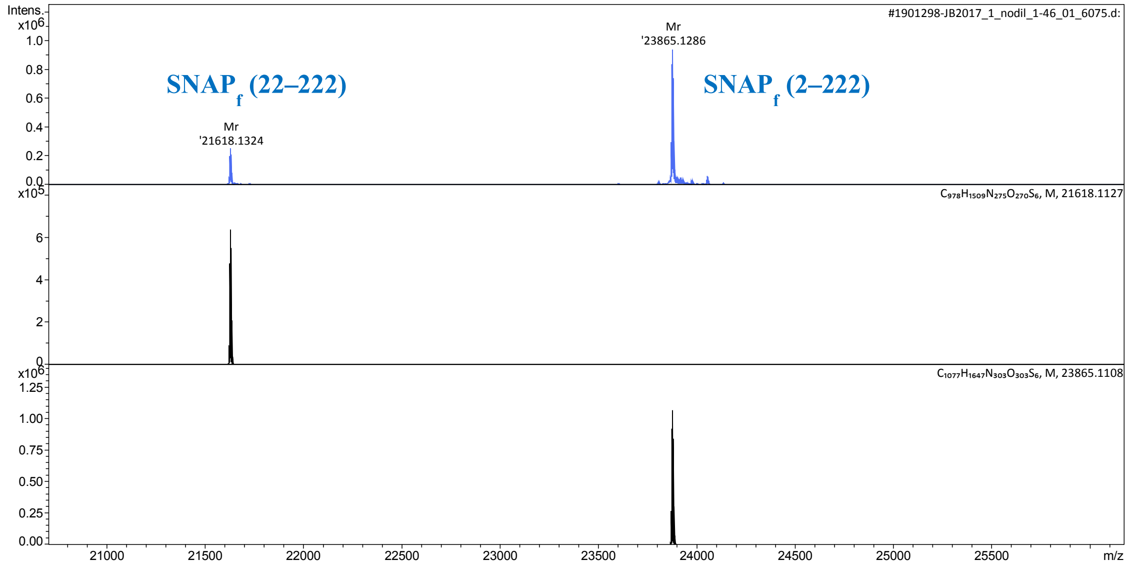


**Supplementary Figure 1. Full protein mass spectrometry on purified SNAP$_f$.** Deconvoluted (upper panel, blue) molecular mass of SNAP$_f$ correlating to two post-translationally processed proteins. Calculated mass of SNAP$_f$ (22-222) (mid panel) and of SNAP$_f$ (2-222) (lower panel) is in agreement with observed masses.

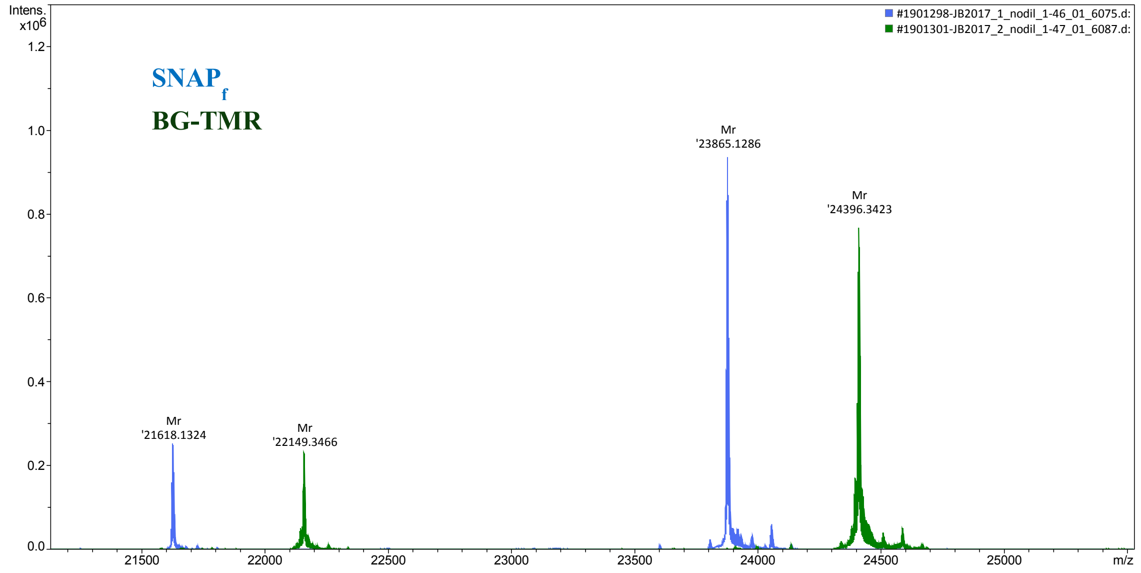


**Supplementary Figure 2. Labeling of SNAP$_f$ by BG-TMR is complete.** Comparison of deconvoluted masses from SNAP$_f$ (blue) and after incubation with BG-TMR (green) shows correct mass shift for labelling and no detectable unlabeled SNAP$_f$.

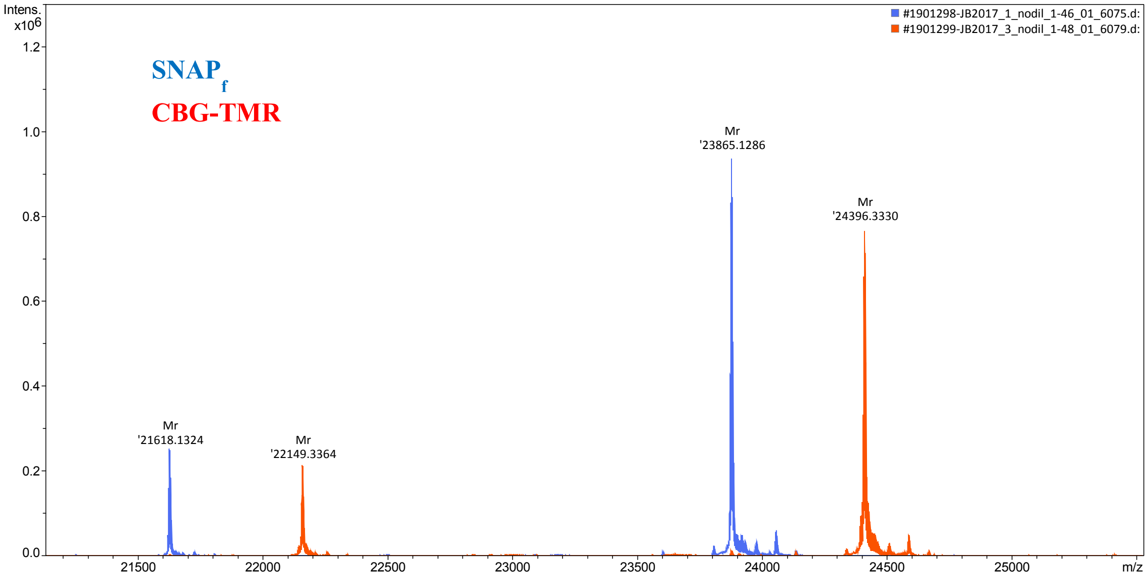


**Supplementary Figure 3. Labeling of SNAPf by CBG-TMR is complete.** Comparison of deconvoluted masses from $SNAP_f$ (blue) and after incubation with CBG-TMR (red) shows correct mass shift for labelling and no detectable unlabeled $SNAP_f$.

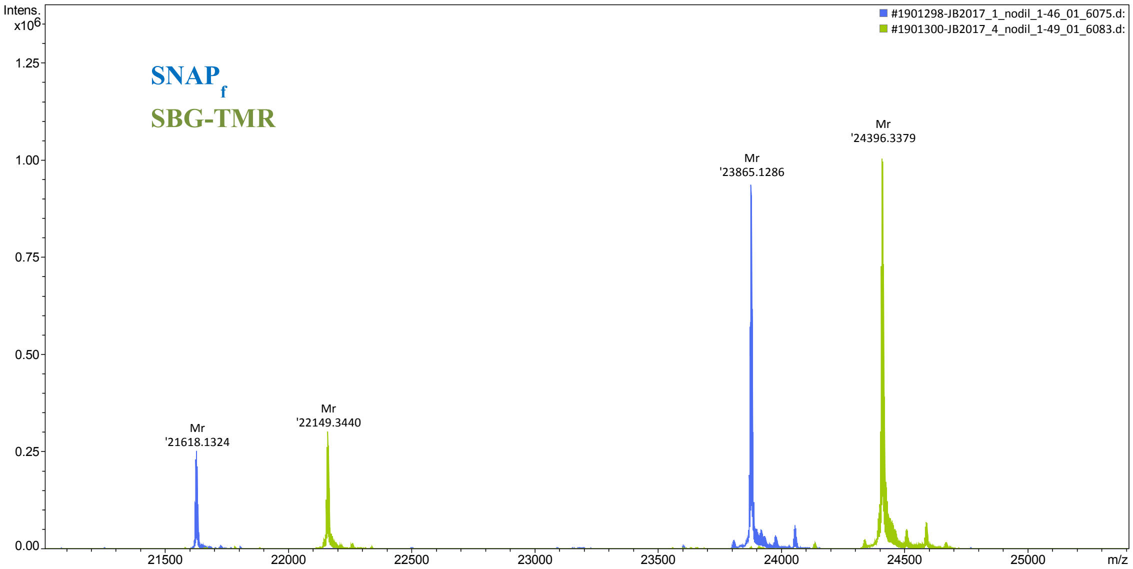


**Supplementary Figure 4. Labeling of SNAPf by SBG-TMR is complete.** Comparison of deconvoluted masses from $SNAP_f$ (blue) and after incubation with SBG-TMR (green) shows correct mass shift for labelling and no detectable unlabeled $SNAP_f$.

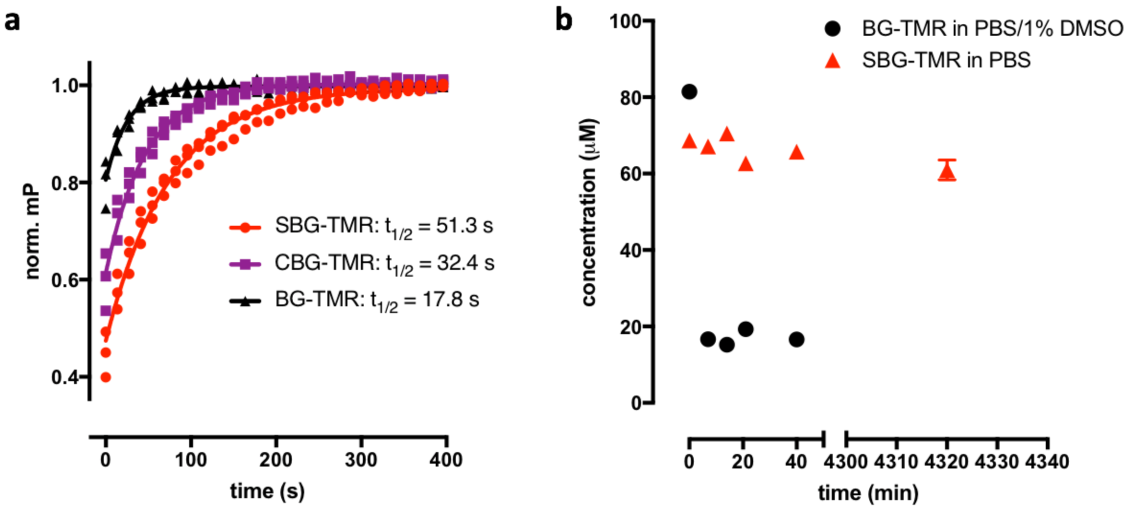


**Supplementary Figure 5. Kinetics and Solubility of TMR-linked SNAP-tag substrates. a**) Kinetics of SNAPf labelling with BG- (black), CBG- (purple) and SBG-TMR (red) by fluorescence polarization. 3 independent measurements, mono-exponential fit. **b**) Solubility of BG-TMR (in PBS + 1% DMSO, black) and SBG-TMR (in PBS, red). While BG-TMR precipitates within minutes, SBG-TMR remains in solution over days.

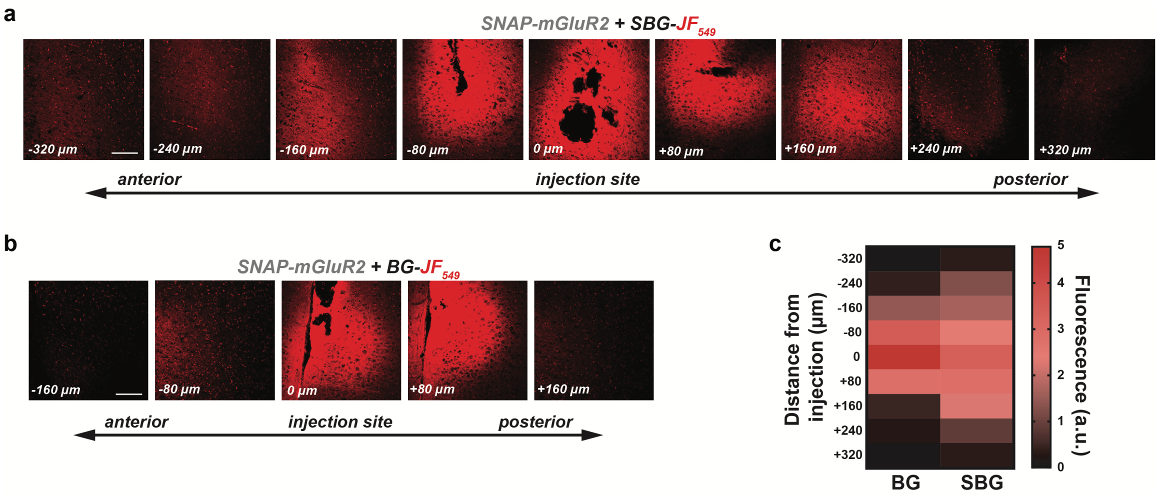


**Supplementary Figure 6. Further analysis of *in vivo* SNAP-tag labeling. a-b,** Images showing SBG-$JF_{549}$ (**a**) and BG-$JF_{549}$ (**b**) labeling in slices anterior and posterior to the injection site. **c,** Summary heat map of fluorescence spread showing wider distribution for SBG compared to BG. Data comes from 3 separate injections for each compound.

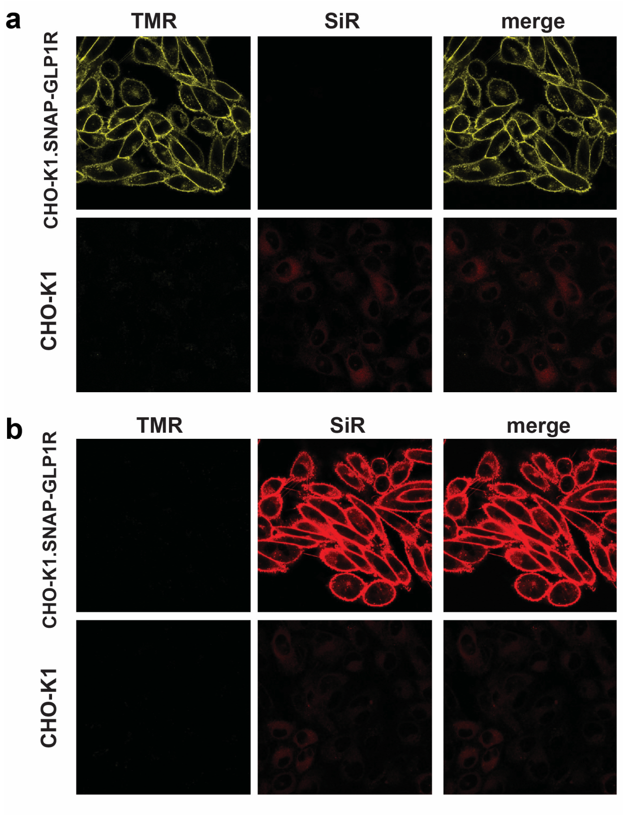


**Supplementary Figure 7. Controls for SNAP-GLP1R labeling conditions. a)** Confocal images of stable SNAP-GLP1R expressing CHO-K1 cells (upper panel) and mock cells (lower panel) labelled with BG-TMR shows no bleedthrough in the SiR channel and absence of labeling in mock cells. **b)** As for (**a**) but with BG-SiR instead of BG-TMR.

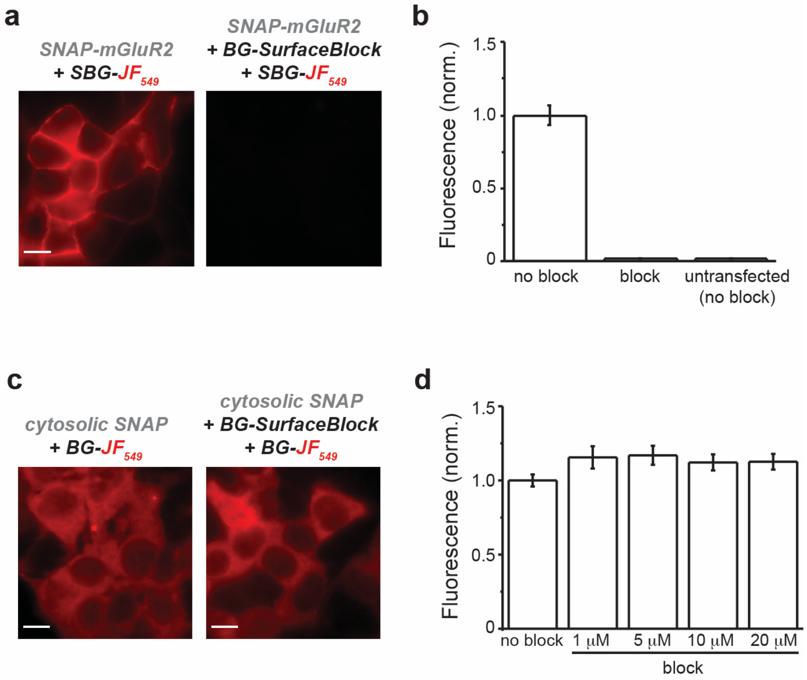


**Supplementary Figure 8. Controls for surface versus intracellular labeling for SIMPull. a-b,** Pre-treatment of HEK 293T cells with BG-surface block prevents labeling of surface SNAP-tags, as seen with representative images (**a**) and summary bar graph (**b**). Following treatment with BG-surface block only background levels (defined by labeling untransfected cells) of SBG-$JF_{549}$ labeling are observed**. c-d**, Pre-treatment with BG-surface block over a wide range of concentrations does not alter labeling efficiency of intracellular SNAP tags as observed in representative images (**c**) or summary bar graph (**d**).
